# Nanoscopy Reveals Heparan Sulfate Clusters as Docking Sites for SARS-CoV-2 Attachment and Entry

**DOI:** 10.1101/2025.09.08.674976

**Authors:** Sue Han, Xin Wang, Tiansheng Li, Ammar Mohseni, Ivan Kosik, Chung Yu Chan, Alberto Domingo López-Muñoz, Jessica Matthias, Reid Suddaby, Zhixiong Wang, Albert J. Jin, Christian A. Wurm, Jonathan W. Yewdell, Ling-Gang Wu

**Author notes:** Equal contribution (Sue Han, Xin Wang, Tiansheng Li).

## Abstract

Virus entry is thought to involve binding a unique receptor for cell attachment and cytosolic entry. For SARS-CoV-2 underlying the COVID-19 pandemic, angiotensin-converting enzyme 2 (ACE2) is widely considered the receptor for cell-surface attachment and cytosolic entry. Using advanced light microscopy to resolve individual virions and receptors, we found instead that heparan sulfate (HS), not ACE2, mediates SARS-CoV-2 cell-surface attachment and subsequent endocytosis. ACE2 functions only downstream of HS to enable viral genome expression. Instead of binding single HS molecules that electrostatically interact with viral surface proteins weakly, SARS-CoV-2 binds clusters of ∼6–137 HS molecules projecting 60–410 nm above the plasma membrane. These tall, HS-rich clusters, present at about one per 6 μm², act as docking sites for viral attachment. Blocking HS binding with the clinically used HS-binding agent pixantrone strongly inhibited an authentic pathogen, the SARS-CoV-2 Omicron JN.1 subvariant, from attaching to and infecting human airway cells. This work establishes a revised entry paradigm in which HS clusters mediate SARS-CoV-2 attachment and endocytosis, with ACE2 acting downstream, thereby identifying HS interactions as a key anti-COVID-19 strategy. This paradigm and its therapeutic implications may apply broadly beyond COVID-19 because, analogous to SARS-CoV-2, HS binds many other viruses but is only considered an attachment regulator.

**Statement of Significance:** Viral entry, a crucial antiviral target, is typically thought to involve binding its unique receptor for the cell surface attachment and subsequent entry. We examined this concept with advanced microscopies to resolve individual receptors and SARS-CoV-2 virions responsible for the COVID-19 pandemic. We discovered two receptors for viral entry: heparan sulfate, a polysaccharide that may bind many viruses, mediates viral attachment and subsequent endocytosis, whereas angiotensin-converting enzyme 2 (ACE2), the generally assumed SARS-CoV-2 receptor, acts only downstream to facilitate viral infection. This model suggests perturbation of HS binding as a more effective anti-COVID-19 strategy than previously recognized. It may apply broadly beyond COVID-19 because, analogous to SARS-CoV-2, HS binds many other viruses but is only considered an attachment regulator.

## Introduction

Understanding how viruses enter host cells is essential for developing effective vaccines and therapies. Viral entry is often initiated by engagement of host receptors and associated co-factors that together facilitate subsequent viral membrane penetration. Decades of research suggest that heparan sulfate, a linear sulfated anionic polysaccharide covalently linked to cell-surface proteoglycans in many cell types, can facilitate attachment of viruses and bacteria by virtue of negative charge interaction with basic virion surface amino acids (*1, 2*). It is thought that viruses exploit weakly interacting with HS to augment the chance of contacting a more specific, high-affinity cell-surface entry receptor (*3*).

A prominent contemporary example of this model is severe acute respiratory syndrome coronavirus 2 (SARS-CoV-2) entry. SARS-CoV-2 is thought to use human angiotensin-converting enzyme 2 (hACE2) as its receptor, based on spike protein (S) high-affinity binding to hACE2, and the requirement of hACE2 for infection (*4–8*). HS binds S *in vitro* enhancing the open conformation of the receptor-binding domain (6, 7). Perturbation of HS significantly reduces SARS-CoV-2 infection in many cell types (*9–11*). These results led to the widely accepted view that SARS-CoV-2 attachment is mediated by SARS-CoV-2 binding to cell surface hACE2, which can be significantly enhanced by the attachment factor HS (*4, 9–11*).

Based on similar experimental results, HS is thought to facilitate the binding of other CoVs to cell surface hACE2 (*2, 3, 12, 13*). This includes SARS-CoV, responsible for the SARS outbreak two decades ago, and human coronavirus NL63, a cause of the common cold (*2, 3, 12, 13*). HS is also considered an attachment factor for medically important non-CoVs including hepatitis C, HIV, Ebola, papilloma, and herpes viruses (*1, 2*). Together, these studies establish HS as an important factor for a broad range of viruses (and other pathogens) that augments pathogen binding with specific cognate cell surface receptors. However, to our knowledge, there is no direct, high-resolution microscopic evidence demonstrating that viruses, including SARS-CoV-2, bind their specific receptors at the cell surface, and not for example, only after internalization into an endosome. Here, we address this decisive lacuna in the currently accepted model for SARS-CoV-2 entry.

To this end, we employed several advanced microscopic methods to achieve unprecedented precision in defining SARS-CoV-2 entry. Using confocal microscopy, stimulated emission depletion microscopy (STED) at ∼60 nm resolution (*14*), and minimal photon flux nanoscopy (MINFLUX) at ∼1-3 nm localization precision (*15, 16*), and electron microscopy (EM), we show that HS, not hACE2 (*4*), is the receptor for SARS-CoV-2 cell-surface attachment and endocytic internalization. Virions exclusively dock at HS clusters that may protrude hundreds of nanometers from the plasma membrane. Internalized virions then bind endosomal hACE2 to deliver viral cores to the cytosol to initiate infection. Blocking HS binding with a clinically used HS-binding agent leads to substantial inhibition of authentic SARS-CoV-2 infection in primary human airway cells. These findings suggest perturbing S-HS binding as a broadly effective strategy for blocking infection by SARS-CoV-2 and other viruses that rely on HS clusters for internalization.

## Results

### Pseudotyped SARS-CoV-2 internalization is independent of ACE2

To observe the virus entering cells and replication, we first used a replicative recombinant vesicular stomatitis virus (VSV) expressing EGFP (infection indicator) and Wuhan-hu-1 SARS-CoV-2 S in place of the natural G receptor/fusion protein (*17, 18*) (VSV*_EGFP_*-S, Fig. 1A). VSV*_EGFP_*-S virus was amplified in Vero or BHK_hACE2_ cells. The amplified VSV*_EGFP_*-S virus was concentrated for purification. The purified VSV*_EGFP_*-S was labeled with Alexa or Atto dyes (e.g., Alexa Fluor 647 NHS ester, abbreviated as A647) via primary amines (*19*) (e.g., VSV*_EGFP_*-S-A647, Fig. 1A, see Methods) for seeing virions entering cells. VSV*_EGFP_*-S-A647 virions attached to glass showed that nearly all viral A647 (V-A647) spots overlapped with immunolabelled S puncta (92.96 ± 0.02%, 606 particles, Fig. 1A). This result suggests that the purified fluorescent V-A647 spots reflect VSV*_EGFP_*-S-A647 virions, rather than non-S-containing fluorescent spots (e.g., VSV*_EGFP_*-A647). We used similar methods to label other recombinant or authentic SARS-CoV-2 virions with various Alexa/Atto dyes (see Methods).

**Fig 1.**
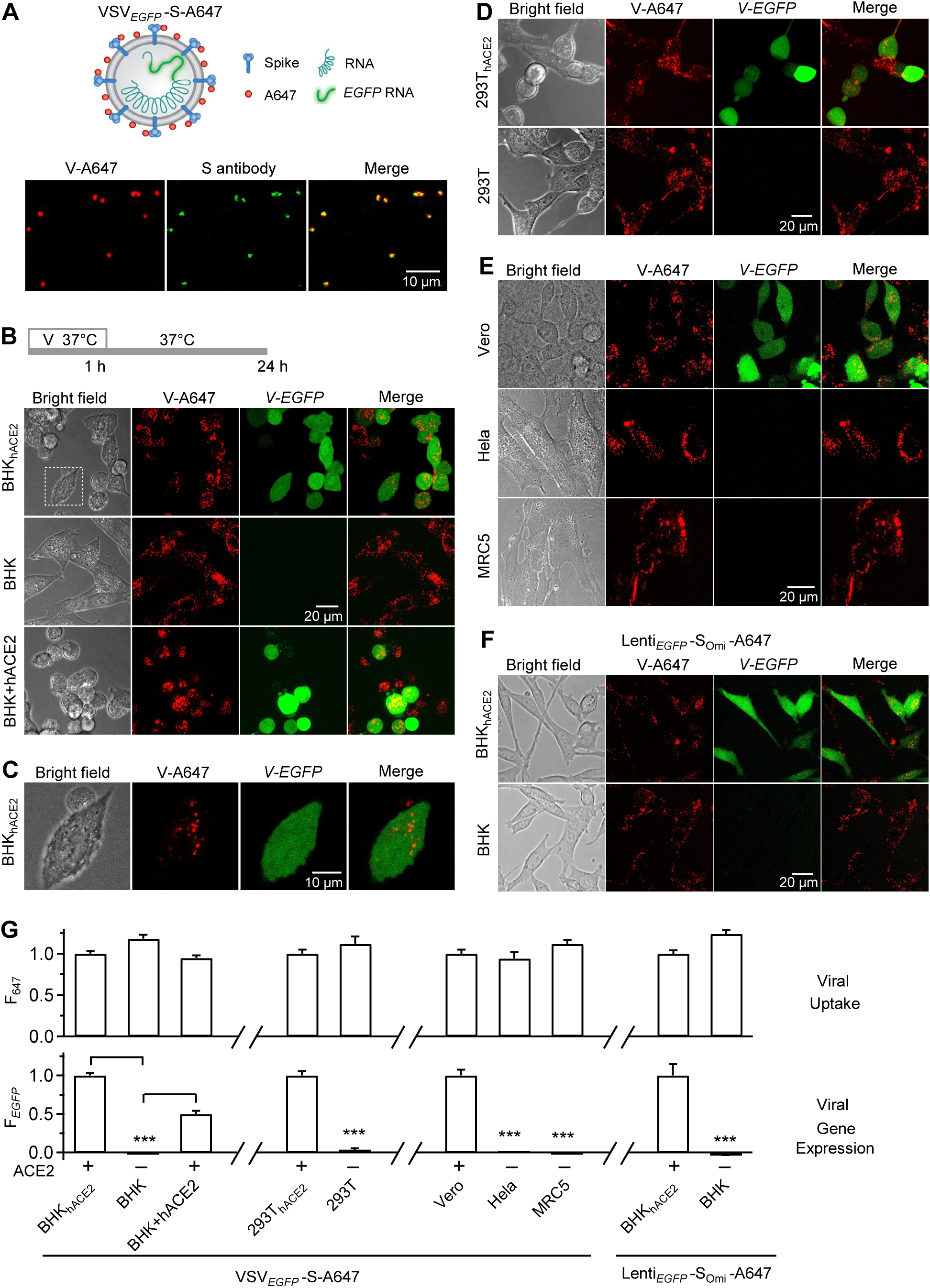
hACE2-independent internalization of pseudo-typed SARS-CoV-2. **A, Upper:** Schematic drawing of a pseudo-typed SARS-CoV-2 virion, VSV*_EGFP_*-S-A647, in which the *EGFP*-coding sequence was inserted into the genomic RNA of VSV, the G protein of VSV was replaced with S of SARS-CoV-2, and A647 was conjugated with primary amines of superficial envelope proteins. **Lower:** Confocal images of A647 of purified VSV*_EGFP_*-S-A647 (V-A647, red) on coverslips and immunofluorescent staining with antibody against S2 subunit of S-protein (green). **B,** Bright-field and confocal images of VSV*_EGFP_*-S-A647’s A647 (V-A647) and *EGFP* expression (*V-EGFP*) in BHK21 cells stably expressing hACE2 (BHK_hACE2_), BHK21 cells without ACE2 expression (BHK), and BHK cells transiently overexpressed with hACE2-BFP plasmid (BHK+hACE2). Top: Incubation protocol – cells were incubated with VSV*_EGFP_*-S-A647 for 1 h at 37°C, followed by washout for 24 h (applies to panels B-G). **C,** The white dotted box in panel B enlarged to show V-A647 spots in EGFP-labelled cytosol. **D,** Similar to panel B, except in 293T_hACE2_ and 293T cells. **E,** Similar to panel B, except in Vero (containing ACE2), Hela (no ACE2) and MRC5 (no ACE2) cells. **F,** Similar to panel B, except VSV*_EGFP_*-S-A647 was replaced with Lenti*_EGFP_*-S_Omi_-A647. **G,** Fluorescence intensity of VSV*_EGFP_*-S-A647’s V-A647 (F_647_) and *V-EGFP* (F*_EGFP_*) in BHK_hACE2_ (471 cells, 3 independent experiments), BHK (370 cells, 3 independent experiments), BHK+hACE2 (515 cells, 3 independent experiments). 293T_hACE2_ (404 cells, 3 independent experiments), 293T (234 cells, 3 independent experiments), Vero (295 cells, 3 independent experiments), Hela (305 cells, 3 independent experiments), and MRC5 (237 cells, 3 independent experiments) cells. Lenti*_EGFP_*-S_Omi_-A647’s F_647_ and F*_EGFP_* in BHK_hACE2_ (364 cells, 3 independent experiments) and BHK (335 cells, 3 independent experiments) cells are also plotted. Virus incubation protocol is shown in panel B. F_647_: mean ± s.e.m., reflecting viral uptake; F*_EGFP_*: mean ± s.e.m., reflecting viral infection. F_647_ and F*_EGFP_* were measured per cell identified in the bright field and were normalized to cells expressing ACE2 in each subgroup (see gaps between subgroups). ***: p < 0.001, t-test.

4-24 h after 1 h VSV*_EGFP_*-S-A647 incubation with cells at 37°C (protocol in Fig. 1B), viral *EGFP* (*V-EGFP*) expression, which indicates viral genome expression, was observed in various ACE2-expressing cells, including hACE2-stably-expressed BHK21 (BHK_hACE2_) or 293T cells (293T_hACE2_), and Vero cells expressing ACE2 endogenously; in contrast, *V-EGFP* was not expressed in non-ACE2-expressing cells, including BHK21 (BHK), 293T, Hela and MRC5 cells (Figs. 1B-G and S1). Similarly, flow cytometric measurements of V-EGFP fluorescence from many thousand cells showed *V-EGFP* expression in BHK_hACE2_ but not BHK cells (Fig. S2). Transient hACE2 overexpression restored *V-EGFP* expression in BHK cells (Figs. 1B-lower and 1G). These results confirmed the hACE2-dependence of SARS-CoV-2 infection (*6, 7, 20–22*).

Surprisingly, V-A647 spot intensity and distribution were independent of ACE2 in the seven cell lines tested (Figs. 1B-E, summarized in Figs. 1G and S1). Confocal sectioning revealed that these V-A647 spots were in the cytoplasm (Figs. 1C and S3), suggesting that VSV*_EGFP_*-S-A647 internalization is independent of ACE2.

Extending these observations, incubating BHK_hACE2_ or BHK cells with Lenti*_EGFP_*-S_Omi_-A647 an A647-labeled, *EGFP*-expressing lentivirus (Lenti*_EGFP_*) pseudotyped with S-protein Omicron variant (S_omi_), revealed similar hACE2-dependent *EGFP* expression, but hACE2-independent virion internalization (Figs. 1F, 1G and S1). While showing that VSV-S and Lenti-S_Omi_ infection is hACE2-dependent as expected, these results revealed surprisingly that VSV-S and Lenti-S_Omi_ internalization is hACE2-independent.

### Endocytic internalization is the main viral infection pathway

Given that hACE2 is required for the fusogenic activity of pseudotyped or authentic SARS-CoV-2 (*4*), the hACE2-independent internalization of V-A647 spots from VSV*_EGFP_*-S-A647 or Lenti*_EGFP_*-S_Omi_-A647 (Fig. 1) may reflect endocytic internalization. Four lines of evidence described below strengthen this suggestion.

First, to track endocytic vesicles, we included a fluorescent dye, Atto 490LS (A490), in the bath solution, which can be taken up by endocytosis into, and thus label, the endocytic vesicle (Figs. S3A and 2A) (*23, 24*). BHK_hACE2_ or BHK cells were incubated with VSV*_EGFP_*-S-A647 plus A490 in the bath solution for 1 h at 37°C, followed by washout. 24 h later, we found near complete colocalization of cytoplasmic V-A647 with A490 spots in either BHK_hACE2_ or BHK cells (Fig. S3A-C). This result suggests that endocytosis takes up both the virus and the bath A490 in an ACE2-independent manner.

Second, we adsorbed VSV*_EGFP_*-S-A647 to BHK_hACE2_ or BHK cells for 1 h at 4°C to prevent endocytosis (cold temperature prevents endocytosis) and then incubated cells with A490 at 37°C for 3 min to allow for, and thus image, endocytic vesicle formation within 3 min (Fig. 2A). STED microscopy showed that most V-A647 spots contained no A490, but colocalized with the plasma membrane (PM) (Fig. S4, 14 cells, 4 cultures) and thus labeled the cell outline. Inside this outline, V-A647 spots colocalized with A490 spots in either BHK_hACE2_ or BHK cells, indicating ACE2-independent endocytic uptake of the virus with the bath A490 as early as 3 min (11 cells, 3 independent experiments; Fig. 2A-C). Consistent with this observation, prolonged washout (4-24 h) completely depleted V-A647 at the cell-surface outline while increasing cytoplasmic V-A647 (Fig. 1B-F), suggesting that surface-attached virions are taken up by endocytosis within 4 h.

**Fig 2.**
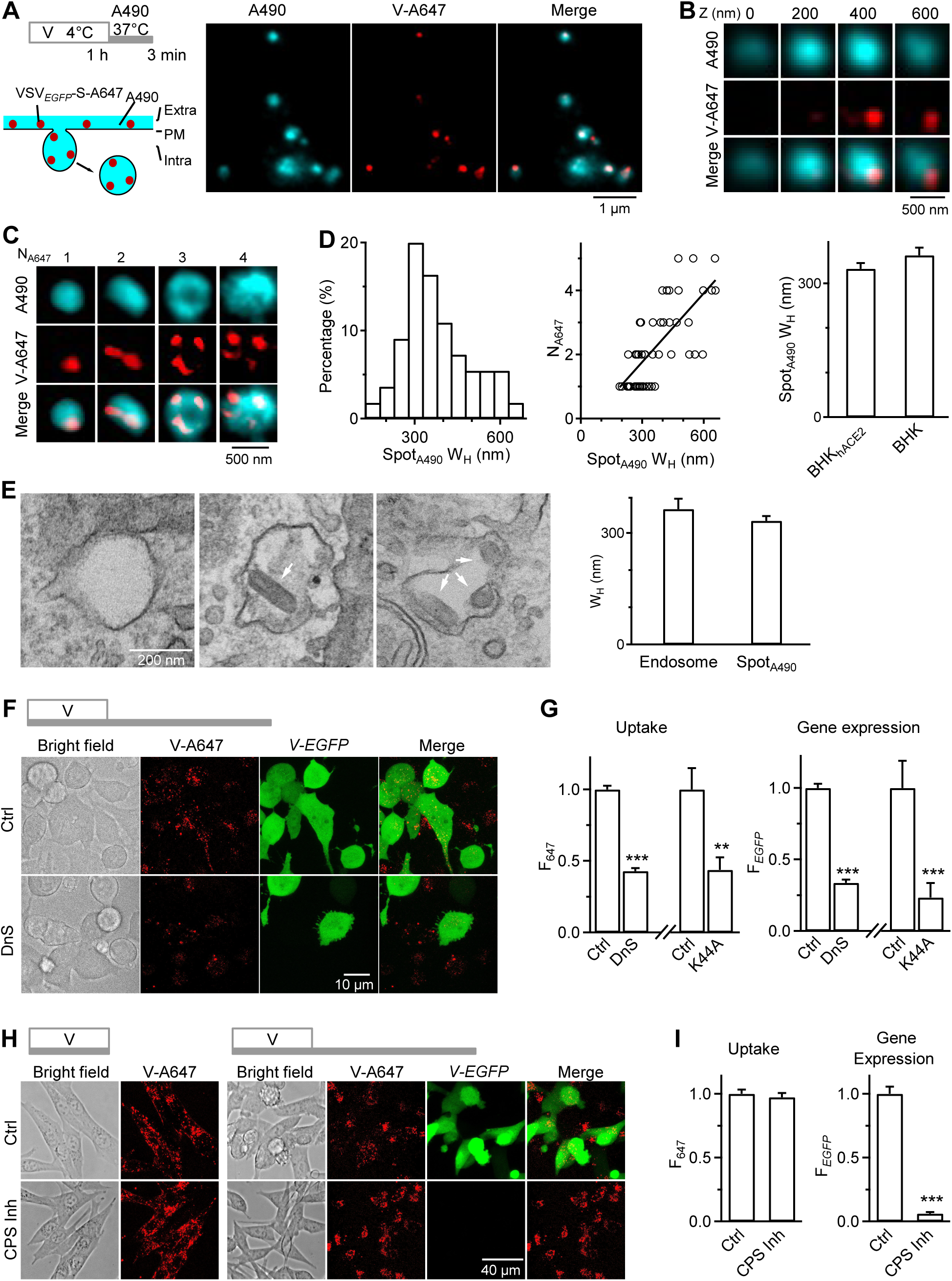
Endocytosis is the primary route for pseudo-typed SARS-CoV-2 internalization and gene expression. **A, Left upper:** incubation protocol for viral attachment and early endocytosis – incubating cells with VSV*_EGFP_*-S-A647 for 1 h at 4°C, followed by 3 min washout with 500-750 μM A490 at 37°C (applies to panel A-D). **Left lower:** strategy for labeling endocytic vesicles taking up PM-attached virions (VSV*_EGFP_*-S-A647, red) with a fluorescent dye in the bath (A490, cyan). **Right:** STED images showing V-A647 spots (red dots) colocalization with A490 spots (cyan) in the cytoplasm (cell boundary beyond the plot). Representative images are from BHK cells. **B**, STED XY-plane images of V-A647 and A490 at different Z sections as labeled showing a cytosolic A490 spot (endocytic vesicle) containing a single V-A647 spot that reflects a single VSV-S virion. Representative images are from BHK cells. **C,** STED XY-plane images of V-A647 and A490 showing a cytosolic A490 spot may contain 1-4 V-A647 spots, each reflecting a single virion. Top: the V-A647 spot number (N_A647_) is labeled. Representative images from BHK cells. **D, Left:** W_H_ distribution of A490 spots (Spot_A490_) containing V-A647 spots (55 Spot_A490_, 3 independent experiments from BHK cells). **Middle:** N_A647_ plotted versus the corresponding Spot_A490_ W_H_ (55 Spot_A490_, 3 independent experiments, from BHK cells). A linear regression fit (correlation coefficient: 0.53) showed that larger Spot_A490_ contains more V-A647 spots. **Right:** V-A647-containing Spot_A490_ W_H_ in BHK_hACE2_ (70 Spot_A490_, 3 independent experiments) and BHK cells (55 Spot_A490_, 3 independent experiments). **E, Left:** EM images showing that endosomes may contain one or more VSV*_EGFP_*-S-A647 virions (arrows). Representative images are from BHK_hACE2_ cells. Incubation protocol: VSV*_EGFP_*-S-A647 for 1 h at 4°C, 15 min washout at 37°C. **Right:** W_H_ (mean ± s.e.m.) of EM-observed virus (VSV*_EGFP_*-S-A647)-containing endosome (38 endosomes, 2 independent experiments) and STED-observed virus (VSV*_EGFP_*-S-A647)-containing Spot_A490_ (70 Spot_A490_, 3 independent experiments). **F,** Bright field and confocal images of V-A647 (uptake) and *V-EGFP* (viral gene expression) in BHK_hACE2_ cells in control (Ctrl) or treated with dynasore (DnS, 80 μM, bath). Top: VSV*_EGFP_*-S-A647 incubation protocol – 1 h incubation, 24 h washout, all at 37°C. **G,** VSV*_EGFP_*-S-A647’s F_647_ (uptake) and F*_EGFP_* (gene expression) in cells in Ctrl (789 cells, 3 independent experiments), treated with DnS (653 cells, 3 independent experiments), or overexpressed with dynamin 2 K44A (19 cells, 2 independent experiments). Virus incubation protocol shown in panel F. **: p < 0.01; ***: p < 0.001, t-test. **H, Left:** bright field and confocal images of V-A647 (uptake) in BHK_hACE2_ cells in Ctrl or treated with cathepsin inhibitor 1 (CPS Inh, 10 μM). Top: VSV*_EGFP_*-S-A647 incubation protocol – 1 h incubation at 37°C. **Right:** bright field and confocal images of V-A647 (uptake) and *V-EGFP* (gene expression) in BHK_hACE2_ cells in Ctrl or treated with CPS Inh. Top: VSV*_EGFP_*-S-A647 incubation protocol – 1 h incubation, 24 h washout, all at 37°C. **I,** F_647_ (uptake) or F*_EGFP_* (gene expression) in cells in Ctrl (540 cells, 3 independent experiments) or treated with CPS Inh (742 cells, 3 independent experiments). F_647_ and F*_EGFP_* were measured after VSV*_EGFP_*-S-A647 incubation in panel H-left and H-right, respectively. ***: p < 0.001, t-test.

Third, 3D-STED imaging showed that A490-labeled vesicle’s full-width-at-half-maximum (W_H_) was 363 ± 17 nm (n = 55) in BHK cells, similar to that (333 ± 13 nm, n = 70) in BHK_hACE2_ cells (Fig. 2B-D), supporting a similar endocytic path regardless of hACE2 presence or not. As W_H_ increased, V-A647 spot number per vesicle increased from 1 to 5 (Figs. 2B and 2C). V-A647 spots appeared cylinder-shaped (length: 135-224 nm) or round-shaped (W_H_: 74-118 nm) (Figs. 2B and 2C). The cylinder shape is consistent with ∼180×70 nm bullet-shaped VSV virions (*19, 25*), whereas the round shape may reflect the cross-section of the cylinder shape.

Thin-section (70 nm) EM showed endosome-like structures with a diameter of 365 ± 28 nm (Range: 115 - 789 nm; 38 structures) containing 1-3 virions with the expected width and length of VSV (*19, 25*) (Fig. 2E). The EM-measured diameter of endosomes containing virions was similar to the STED-measured W_H_ of A490-labeled vesicles containing V-A647 spots (Fig. 2E), strengthening the STED observation of endocytic vesicles. The EM-observed maximal virion number (3) in an endosome was less than the STED-observed (5), likely because we counted virions only in one thin section with EM, but the entire A490-labeled vesicle with 3D-STED microscopy.

The large size (115 - 789 nm) of the virus-containing vesicles observed with STED and EM implies macropinocytosis that may form large vesicles engulfing multiple virions. Our observation is consistent with a recent study showing that a macropinocytosis inhibitor 5-[N-ethyl-N-isopropyl] amiloride (EIPA) (*26*) reduces SARS-CoV-2 replication in Vero cells (*27*).

Fourth, dynasore or dominant-negative dynamin 2-K44A overexpression, which inhibits fission of dynamin-dependent endocytosis (*28–30*), substantially reduced V-A647 internalized 1-24 h after viral incubation (Figs. 2F-G, S5). This result suggests that viruses are internalized via dynamin-dependent endocytosis.

In addition to inhibiting V-A647 endocytosis, dynasore or dynamin 2-K44A inhibited *V-EGFP* expression 8-24 h after virus incubation by ∼66-77% (Figs. 2F-G, S5), suggesting that endocytosis is the main route for viral genome expression. Further strengthening this suggestion, cathepsin inhibitor 1, which inhibits endosomal cathepsin L crucial for SARS-CoV-2 endosomal fusion and infection (*31–34*), nearly abolished *V-EGFP* expression without affecting V-A647 uptake (Fig. 2H and 2I). We concluded that endocytic internalization is the major pathway for pseudotyped SARS-CoV-2 genome expression.

### Heparan sulfate, but not hACE2, is required for viral cell-surface attachment and internalization

Four sets of evidence suggest that heparan sulfate (HS), but not hACE2, is the receptor underlying VSV-S cell-surface attachment. First, VSV-S and Lenti-S_Omi_ internalization is hACE2-independent (Figs. 1 and S1), implying that the cell-surface attachment of these viruses is independent of hACE2. Second, with 1 h viral incubation at 4°C for virus attachment at the cell surface, V-A647 intensity at the cell surface was similar between BHK_hACE2_ cells that contained hACE2 at the cell surface and BHK cells that did not contain hACE2 (Figs. 3A and 3B), revealing hACE2-independent viral cell-surface attachment. Further supporting this conclusion, flow cytometric measurements from many thousands of cells showed similar V-A647 intensity between BHK_hACE2_ and BHK cells after 1 h viral incubation at 4°C for viral cell-surface attachment (Fig. S6). Flow cytometry also showed that the V-A647 attachment at the cell surface of BHK cells was similar to that in BHK_hACE2_ cells over a ∼600-fold dilution of the virus concentration (Fig. S7), indicating that the virus cell-surface attachment is independent of hACE2 across a wide range of virus concentrations. Third, 93-94% V-A647 spots colocalized with immunolabelled HS spots in BHK_hACE2_ (94 ± 2%, 8 cells) and BHK (93 ± 2%, 8 cells) cells (Fig. 3C), indicating hACE2-independent VSV-S colocalization with HS. HS spots may thus serve as the docking site for VSV-S attachment at the cell surface. Fourth, heparinase I/II/III mixture (HPRase), which nearly abolished cell-surface immunolabelled HS (Fig. 3D) (*35*), reduced cell-surface V-A647 by ∼83-85% in both BHK_hACE2_ and BHK cells (after 1 h viral incubation at 4°C for viral cell-surface attachment, Fig. 3E), suggesting that HS crucial for VSV-S cell-surface attachment. Application of HPRase also reduced intracellular V-A647 spots by ∼81-82% in both BHK_hACE2_ and BHK cells, suggesting that HS-mediated viral attachment is a critical step preceding viral endocytosis (Fig. 3F; incubation protocol: 1 h virus incubation at 4°C for viral attachment, followed by 1 h washout at 37°C for viral uptake). Further, HPRase reduced *V-EGFP* expression in BHK_hACE2_ cells by ∼79%, suggesting that HS-mediated viral attachment is a critical step preceding viral genome expression (24 h after virus incubation, Fig. 3G). In summary, these results (Fig. 3A-G) suggest that HS binding with VSV-S underlies viral cell-surface attachment, which is crucial for the subsequent virus endocytosis and genome expression.

**Fig 3.**
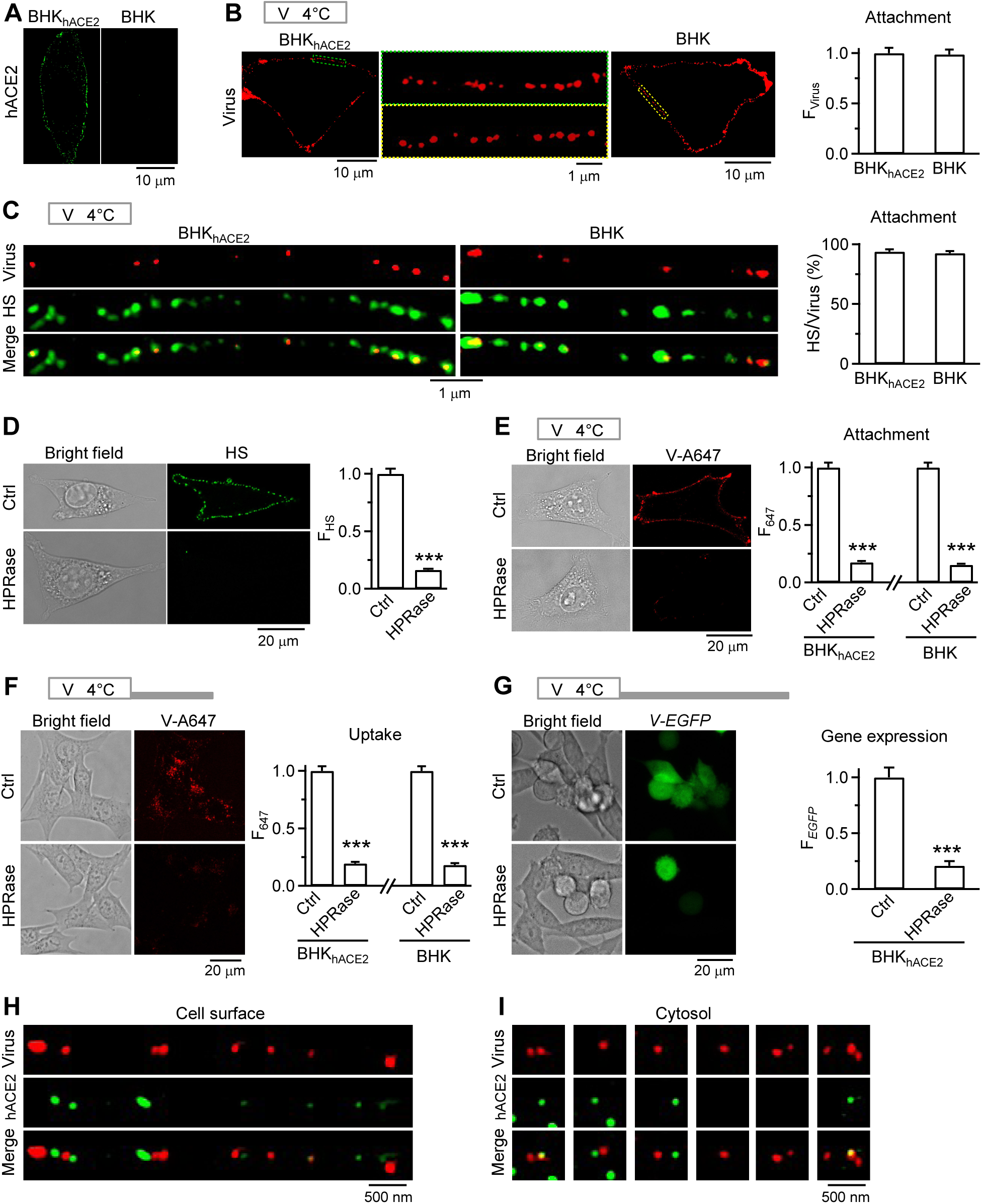
Heparan sulfate is the receptor for pseudo-typed SARS-CoV-2 attachment at the cell surface. **A,** Immunolabelling (A647) of hACE2 in a BHK_hACE2_ (left) and a BHK (right) cell. **B,** ACE2-independent viral cell-surface attachment. **Images:** STED images showing virus cell-surface attachment in a BHK_hACE2_ (left) or BHK (right) cell. Middle: images in green (left) and yellow (right) boxes replotted on a larger scale. Top: incubation protocol – virus (VSV*_EGFP_*-S-A490) 1 h at 4°C for viral attachment at the cell surface (applies to panels B, C, E, and H). **Bar graph:** Cell-surface virus fluorescence intensity (F_Virus_) per BHK_hACE2_ (92 cells, 3 independent experiments) or BHK cell (98 cells, 3 independent experiments). Data normalized to the mean of the BHK_hACE2_ group. **C,** Cell-surface-attached viruses colocalized with HS puncta. **Images:** STED images of viral spots (VSV*_EGFP_*-S-A490’s V-A490) and immunolabelled HS (Alexa 488-conjugated antibody) at BHK_hACE2_ or BHK cell surface. Top: incubation protocol – VSV*_EGFP_*-S-A647 1 h at 4°C for cell-surface attachment. **Bar graph:** the percentage of viral spots overlapping with HS puncta in BHK_hACE2_ (8 cells, 4 independent experiments) or BHK cells (8 cells, 4 independent experiments). **D,** Bright field and confocal images of immunolabelled HS (left) and HS fluorescence intensity (F_HS_, right) in BHK_hACE2_ cells in Ctrl (188 cells, 3 independent experiments) or treated with Heparinases I/II/III (HPRase, 264 cells, 3 independent experiments). ***: p < 0.001, t-test. **E, Images:** bright field and confocal image of V-A647 in BHK_hACE2_ cells in Ctrl or treated with HPRase. Top: incubation protocol – VSV*_EGFP_*-S-A647 1 h at 4°C for cell-surface attachment. **Bar graph:** cell-surface F_647_ per cell for BHK_hACE2_ cells (left) in Ctrl (125 cells, 3 independent experiments) or treated with HPRase (150 cells, 3 independent experiments), and for BHK cells (right) in Ctrl (147 cells, 3 independent experiments) or treated with HPRase (127 cells, 3 independent experiments). ***: p < 0.001, t-test. **F, Images:** bright field and confocal image of V-A647 (uptake) in BHK_hACE2_ cells in Ctrl or treated with HPRase. Top: incubation protocol – VSV*_EGFP_*-S-A647 1 h at 4°C (for viral attachment), 1 h washout at 37°C (for endocytosis of cell-surface-attached viruses). ***: p < 0.001, t-test. **Bar graph:** F_647_ (uptake) for BHK_hACE2_ cells in Ctrl (201 cells, 3 independent experiments) or treated with HPRase (197 cells, 3 independent experiments), and for BHK cells in Ctrl (202 cells, 3 independent experiments) or treated with HPRase (206 cells, 3 independent experiments). **G, Images:** bright field and confocal image of *V-EGFP* expression in BHK_hACE2_ cells in control or treated with HPRase. Top: incubation protocol – VSV*_EGFP_*-S-A647 1 h at 4°C, 24 h washout at 37°C. ***: p < 0.001, t-test. **Bar graph:** F_EGFP_ (viral gene expression) per BHK_hACE2_ cell in Ctrl (482 cells, 3 independent experiments) or treated with HPRase (391 cells, 3 independent experiments). **H,** STED images of viral spots (VSV*_EGFP_*-S-A490’s A490) and immunolabelled hACE2 puncta (A647) at a BHK_hACE2_ cell surface. **I,** STED images of viral spots (VSV*_EGFP_*-S-A490’s A490) and immunolabelled hACE2 puncta (A647) inside BHK_hACE2_ cells – hACE2 is sometimes taken up with viruses in endosomes.

### hACE2 acts downstream of endocytosis to promote viral infection

We showed that hACE2 is essential for viral genome expression but not cell-surface viral attachment or endocytosis (Figs. 1–3), suggesting that hACE2 acts downstream of endocytosis. Unlike HS spots, only 22 ± 2% cell-surface V-A647 spots colocalized with hACE2 spots in BHK_hACE2_ cells (Fig. 3H; 13 cells, 2 independent experiments). However, 38 ± 3% cell-surface V-A647 spots were within 200 nm from neighboring hACE2 spots (Fig. 3H, 13 BHK_hACE2_ cells, 2 independent experiments). These virions and hACE2 may be packed into the same (large) vesicle, as we noted that 38 ± 5% intracellular V-A647 spots colocalized with or within 200 nm from hACE2 spots (13 BHK_hACE2_ cells, 2 independent experiments; Fig. 3I). These results suggest that hACE2’s infection role may be due to cell-surface hACE2 co-internalization with virions to facilitate cathepsin L-dependent (Fig. 2H-I) (*31–34*) viral fusion at the endosomal membrane.

### VSV-S recapitulates the SARS-CoV-2 entry pathway

To extend our findings to authentic SARS-CoV-2 virions, we repeated the above VSV-S experiments using SARS-CoV-2 Omicron variant (CoV2-S_omi_) (*35, 36*), containing inserted *mCherry* gene and A647- or A490-labeled envelope proteins (CoV2*_mCherry_*-S_Omi_-A647 or CoV2*_mCherry_*-S_Omi_-A490). In addition, we replaced S in VSV-S with S_omi_ for comparison between wild-type (VSV*_EGFP_*-S-A647) and Omicron variants (VSV*_EGFP_*-S_omi_-A647). Three sets of evidence suggest that the authentic SARS-CoV-2 entry path is similar to VSV-S.

First, VSV-S (Figs. 1B-E and 1G), Lenti-S_omi_ (Fig. 1F-G), VSV-S_omi_ (Fig. 4A-B), and CoV2-S_Omi_ (Fig. 4C-D) all exhibited hACE2-independent viral internalization, but hACE2-dependent viral gene (*EGFP* or *mCherry*) expression. Second, similar to VSV-S (Fig. 2A-D), 1-5 CoV2-S_omi_ virions were found in ∼150-700 nm vesicles in the cytoplasm (Fig. 4E, 18 cells, 2 independent experiments). Third, analogous to VSV-S (Fig. 3E), removing HS by HPRase nearly abolished cell-surface-attached CoV2-S_omi_ (Fig. 4F). Based on these observations, we conclude that Omicron variant of SARS-CoV-2 enters BHK cells indistinguishably to VSV-S.

**Fig 4.**
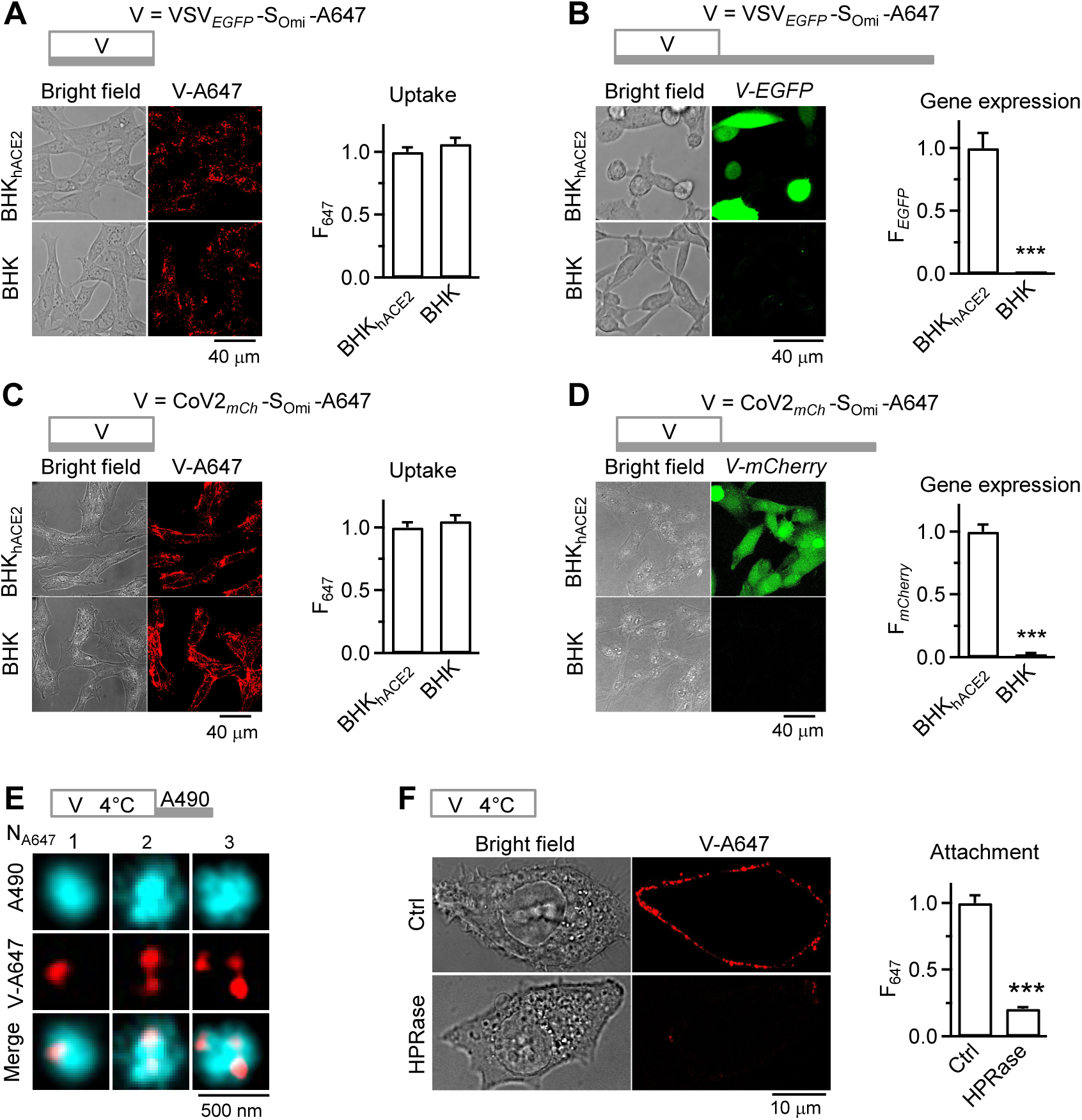
SARS-CoV-2 cell entry path is analogous to pseudo-typed SARS-CoV-2. **A,** Images (bright field and confocal images) of VSV*_EGFP_*-S_omi_-A647’s V-A647 (left) and F_647_ (right, viral uptake) in BHK_hACE2_ (146 cells, 3 independent experiments) or BHK (149 cells, 3 independent experiments) cells. Top: incubation protocol – VSV*_EGFP_*-S_omi_-A647 1 h at 37°C (for virus endocytosis). **B,** Images (bright field and confocal images) of VSV*_EGFP_*-S_omi_-A647’s *V-EGFP* (left) and F*_EGFP_* (right, viral gene expression) in BHK_hACE2_ (273 cells, 3 independent experiments) or BHK (200 cells, 3 independent experiments) cells. Top: incubation protocol – VSV*_EGFP_*-S_omi_-A647 1 h, 24 h washout, all at 37°C (for viral gene expression). **C,** Images (bright field and confocal images) of CoV2*_mCh_*-S_omi_-A647’s A647 fluorescence (V- A647, left) and F_647_ (right, uptake) in BHK_hACE2_ (120 cells, 3 independent experiments) or BHK (120 cells, 3 independent experiments) cells. Top: incubation protocol – CoV2*_mCh_*-S_omi_-A647 1 h at 37°C (for virus endocytosis). **D,** Images (bright field and confocal images) of CoV2*_mCh_*-S_omi_-A647’s *mCherry* expression (*V-mCherry*, pseudo color, left) and F*_mCherry_* (right, uptake) in BHK_hACE2_ (120 cells, 3 independent experiments) or BHK (120 cells, 3 independent experiments) cells. Top: incubation protocol – CoV2*_mCh_*-S_omi_-A647 1 h, 6 h washout, all at 37°C (for viral gene expression). **E,** STED XY-plane images of CoV2*_mCh_*-S_omi_-A647’s V-A647 and A490 in cytosol showing that the number (N_A647_) of V-A647 spots (virions) in a A490 spot (endocytic vesicle) may vary as labelled. Top: incubation protocol – CoV2*_mCh_*-S_omi_-A647 1 h at 4°C, 3 min washout with 500-750 μM A490 at 37°C. **F,** Images (bright field and confocal images) of CoV2*_mCh_*-S_omi_-A647 ’s V-A647 (left) and surface F_647_ per cell (right) for BHK_hACE2_ cells in Ctrl (120 cells, 3 independent experiments) or treated with HPRase (120 cells, 3 independent experiments). Top: incubation protocol – CoV2*_mCh_*-S_omi_-A647 1 h at 4°C for cell-surface attachment.

### Structural arrangement of HS as the SARS-CoV-2 attachment receptor

We characterized HS topology at molecular resolution to better understand its function as the virus receptor. 3D confocal imaging of immunolabelled HS revealed 563 ± 38 HS-spots per BHK_hACE2_ cell-surface (range: 244-860 spots, 16 cells, 4 independent experiments, Fig. 5A). Given an average cell-surface area of 3412 ± 360 μm^2^ (14 cells, Fig. S8), this amounts to an average of 1 HS-spot per 6 μm^2^ (or 1 per 2.5×2.5 μm^2^). 2-D MINFLUX imaging (∼3-nm localization precision) (*16*) revealed oval-shaped clusters of HS molecules labeled by Flux 647-conjugated anti-HS antibody (HS-Ab-F647); cluster width (parallel to PM) and height (vertical to PM) were 36-243 (133 ± 6 nm) and 63-407 nm (202 ± 10 nm, 53 clusters, 9 cells, Figs. 5B-D and S9A-C). Although individual molecules spread between clusters were also observed (Fig. 5B), MINFLUX-detected HS-clusters corresponded to confocally resolved HS-spots at the PH-mNeonGreen (PH_mNG_)-labeled plasma membrane (Fig. 5B). Virus (CoV2-S_omi_ or VSV-S) incubation resulted in virion colocalization with MINFLUX-detected HS-clusters (Fig. 5C) or confocally resolved HS-spots (Fig. 3C), indicating that SARS-CoV-2 virions dock at HS-clusters rather than individual HS molecule alone.

**Fig 5.**
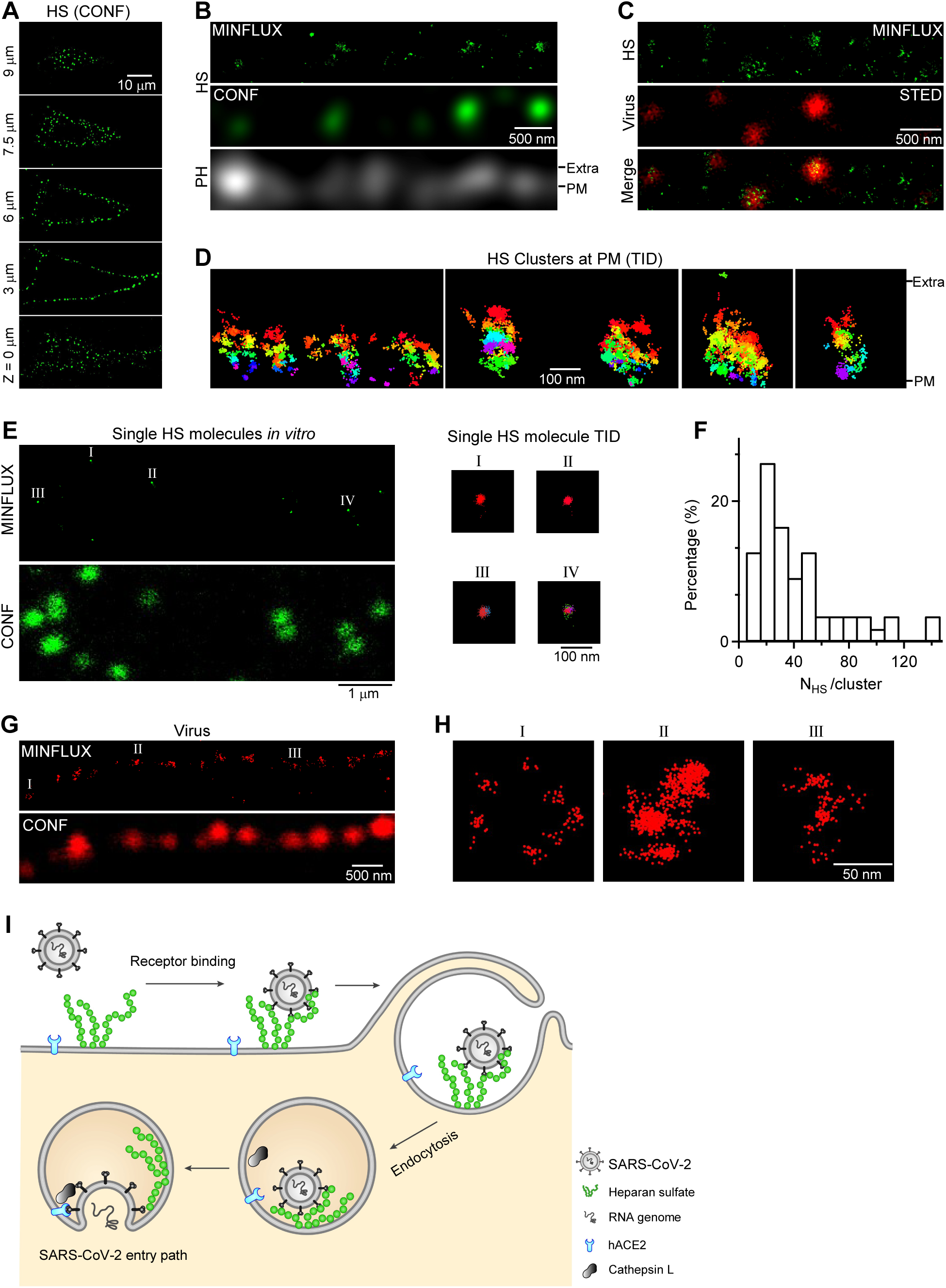
Structural arrangement of HS as the viral receptor at the cell surface. **A,** Confocal (CONF) XY-plane images of immunolabelled HS (A647-conjugated secondary antibody) at various Z-axis locations as labeled (16 cells, 4 independent experiments). **B,** MINFLUX (2D) pixel-based rendering of immunolabelled HS (HS-Ab-F647, see Methods), the corresponding confocal (CONF) image of immunolabelled HS, and confocal PH_mNG_ (labeling PM) image at a BHK_hACE2_ cell surface [extracellular (Extra) compartment is above the PM]. **C,** MINFLUX image of immunolabelled HS and STED image of CoV2*_mCh_*-S_omi_-A490 (Virus) at a BHK_hACE2_ cell surface (merged at the bottom). Cells were incubated with CoV2*_mCh_*-S_omi_-A490 for 1 h at 4°C for viral cell-surface attachment. Top is the extracellular side. **D,** MINFLUX coordinate-based rendering of HS clusters at the cell surface with each trace ID (TID) labeled in different colors (four different regions are shown). Extra: extracellular side; PM: plasma membrane position. **E, Left:** MINFLUX pixel-based rendering (upper) and confocal image (lower) of immunolabelled (HS-Ab-F647) single HS molecules in vitro (on the coverslip). **Right:** MINFLUX coordinate-based rendering of single HS molecules color-coded by trace IDs (TIDs) (data I, II, III, IV from left panel). **F,** Distribution of HS-molecule number per HS-cluster (N_HS_/cluster, 51 clusters, 8 cells, 3 independent experiments). **G,** MINFLUX pixel-based rendering (upper) and confocal image (lower) of CoV2-S_omi_ (CoV2*_mCh_*-S_omi_-A647’s A647) at the cell surface (extracellular side: above the confocal spots). Cells were incubated with CoV2*_mCh_*-S_omi_-A647 for 1 h at 4°C for viral cell-surface attachment. **H,** MINFLUX coordinate-based rendering of CoV2*_mCh_*-S_omi_-A647 showing three individual virions from panel G (I, II, III in panel G). **I,** SARS-CoV-2 cell-entry model: heparan sulfate molecules in clusters protruding tens to hundreds of nanometers from the plasma membrane serve as the receptor for viral cell-surface attachment; endocytosis is the primary route to internalize surface-attached virions. If endocytosis co-internalizes local hACE2 in the close vicinity to the heparan sulfate cluster at the cell surface, viral RNA genome can be released from endosomes for viral genome expression to infect cells.

The mean diameter [(width+length)/2] of the cell-surface HS-Ab-F647 cluster (168 ± 8 nm, 53 clusters, 9 cells; Fig. S9A, B, see also Fig. 5D) was much larger than that of single HS-Ab-F647 molecule *in vitro* (26 ± 1 nm, 67 molecules, Figs. 5E; S9D and S9E), suggesting that each cell-surface HS cluster is composed of many single HS molecules. Normalizing the number of Flux 647 emission bursts (trace ID) per cell-surface cluster (73 ± 8, from 51 clusters, Fig. S9C) onto that per single molecule (2.0 ± 0.2, 67 molecules, Fig. S9F) yielded 38 ± 4 HS-molecules per cell-surface cluster (range: 6 - 137, 51 clusters, 8 cells, Fig. 5F). The number is likely higher due to steric limitations to antibody binding with individual HS molecules in the cluster.

Importantly, confocally resolved CoV2*_mCh_*-S_omi_-A647 spots corresponded to clusters of MINFLUX-resolved CoV2*_mCh_*-S_omi_-A647’s A647 localizations (Fig. 5G). Many clusters measured ∼70-130 nm in mean diameter (Figs. 5G and 5H), agreeing with SARS-CoV-2 virion diameter of ∼100 nm (*4*); some were larger (Fig. 5G), likely representing multiple virions. These MINFLUX-resolved SARS-CoV-2 images verified our virus labeling strategy (Fig. 1A).

### SARS-CoV-2 JN.1 variant binds to heparan sulfate, not hACE2, in primary human airway cells

To test whether our findings with the engineered Omicron variant (CoV2_mCh_-S_omi_) extend to an authentic strain that infected people, we examined the JN.1 subvariant (BA.2.86.1.1) virion binding in Chinese hamster ovary (CHO) cell lines with glycosaminoglycan (GAG) deficiencies, the CHO-pgs-745 and CHO-pgsB-618 cells, in both of which heparan sulfate biosynthesis is blocked by loss of xylosyltransferase-2 (XYLT2) or β-1,4-galactosyltransferase-7 (B4GALT7) activity (*37–39*). Consistent with our previous observations, Alexa 647-labeled JN.1 variant (CoV2-S_JN.1_-A647) showed robust binding to wild-type CHO-K1 cells, but a ∼81-84% decrease in binding in both CHO-pgsA-745 and CHO-pgsB-618 cells lacking HS (Fig. 6A-B). These results indicate that HS is critical for CoV2-S_JN.1_-A647 attachment to the cell surface.

**Fig 6.**
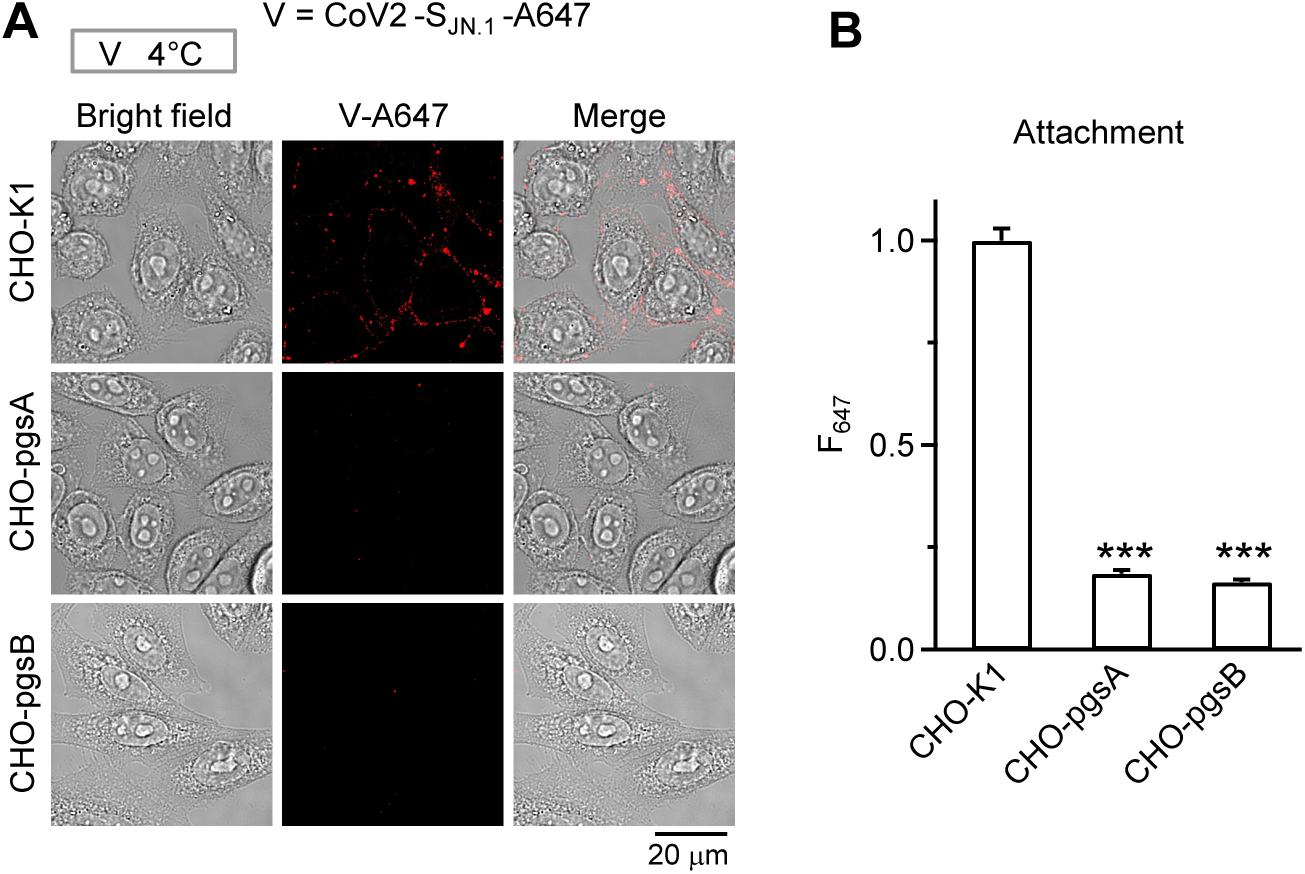
Attachment of SARS-CoV-2 JN.1 variant at the cell surface depends on heparan sulfate. **A,** CoV2-S_JN.1_-A647 attachment at the cell surface: sampled bright field and confocal images of CoV2-S_JN.1_-A647’s V-A647 in wild-type CHO cells (CHO-K1), GAGs-deficient CHO-pgsA-745 cells lacking HS, and GAGs-deficient CHO-pgsB-618 cells lacking HS (images merged in the right) after incubation of CoV2-S_JN.1_ for 1 h at 4°C for labeling cell-surface attachment (incubation protocol shown on the top). **B,** Quantification of CoV2-S_JN.1_-A647 attachment at the cell surface: cell-surface fluorescence of CoV2-S_JN.1_-A647 (F_647_) per cell (mean + s.e.m.) in CHO-K1 (117 cells, 2 repeats), CHO-pgsA-745 (93 cells, 2 repeats), and CHO-pgsB-618 cells (90 cells, 2 repeats). ***: p < 0.001 (ANOVA test).

To further test the clinical relevance, we pre-treated normal human bronchial/tracheal epithelial (NHBE) cells with: 1) control solution, 2) pixantrone (PIX), a drug under clinical trial for cancer treatment due to its DNA intercalating activity, which can bind HS to inhibit HS binding with proteins (*10*), or 3) anti-hACE2 antibody that blocks CoV2-S_JN.1_-A647 binding with ACE2 (*4*). Immunostaining of nucleocapsid protein (N protein) and spike showed that PIX nearly abolished the cell-surface attachment of CoV2-S_JN.1_-A647, while anti-hACE2 antibody had no effect compared to control (Fig. 7A), suggesting that HS, but not ACE2 mediates authentic SARS-CoV-2 cell-surface attachment. Confocal and STED images showed the colocalization of HS and CoV2-S_JN.1_-A647 (Fig. 7B). Analogous to HS colocalization with VSV-S (Fig. 3C) or CoV2*_mCh_*-S_omi_ (Fig. 5C), this result (Fig. 7B) suggests that HS binds CoV2-S_JN.1_-A647, mediating CoV2-S_JN.1_-A647 attachment on the NHBE cell surface.

**Fig 7.**
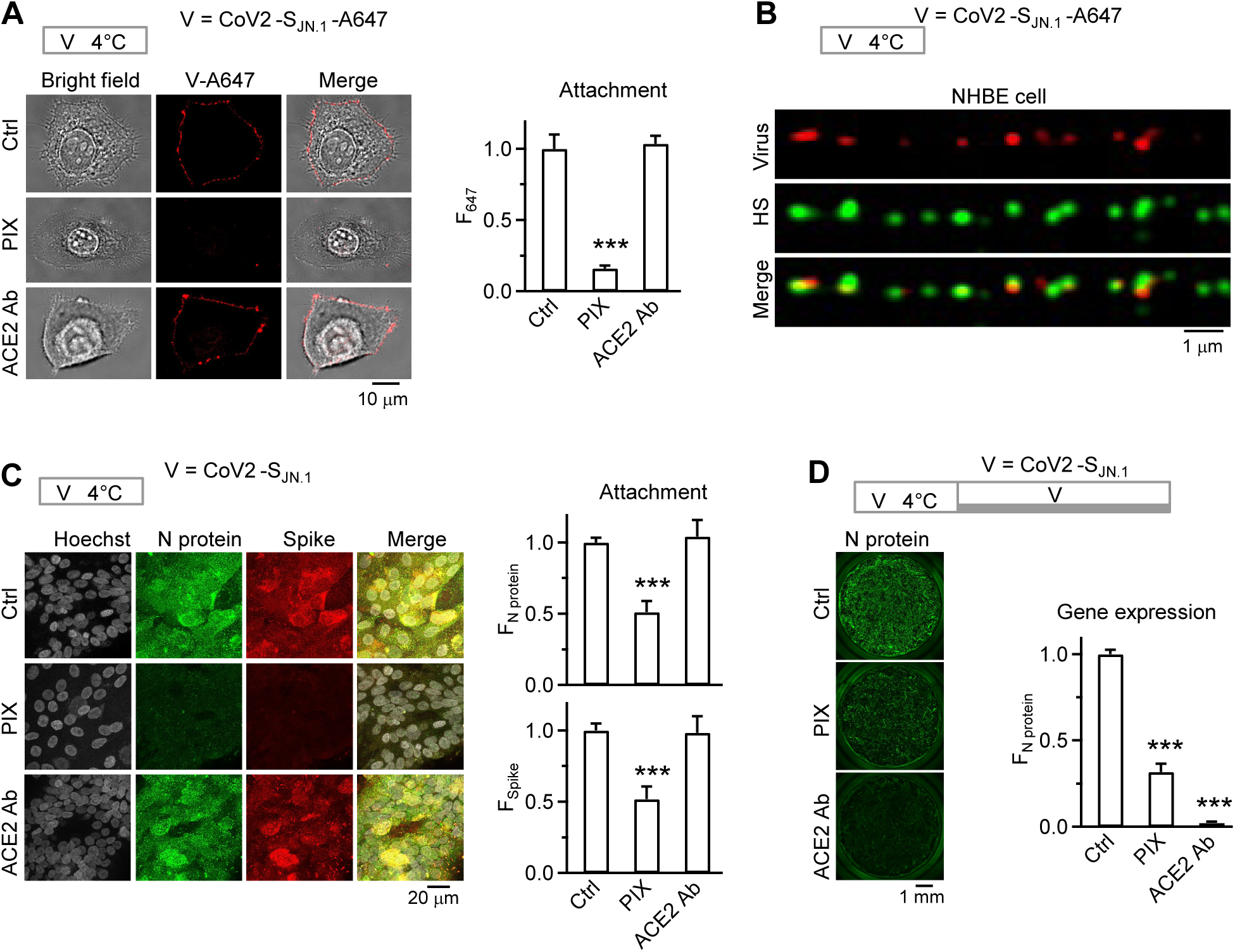
SARS-CoV-2 JN.1 variant attachment and infection in primary human airway cells depend on HS and are inhibited by a clinically used HS-binding agent. **A, Images for virus attachment:** bright field, confocal, and merged images of CoV2-S_JN.1_-A647 in normal human bronchial/tracheal epithelial cells (NHBE) pre-treated with control (Ctrl), pixantrone (PIX), or anti-ACE2 antibody (ACE2 Ab). Top: incubation protocol – CoV2-S_JN.1_-A647 1 h at 4°C for cell-surface attachment. **Bar graph for virus attachment:** cell-surface fluorescence of CoV2-S_JN.1_-A647 (F_647_) per cell for normal human bronchial/tracheal epithelial (NHBE) cells pre-treated with control (Ctrl, 16 cells, 2 independent experiments), pixantrone (PIX, 18 cells, 2 independent experiments), or anti-ACE2 antibody (ACE2 Ab, 24 cells, 2 independent experiments). ***: p < 0.001 (ANOVA test). **B,** Cell-surface-attached viruses colocalized with HS puncta: confocal images of viral spots (CoV2-S_JN.1_-A647’s V-647, red) and immunolabelled HS (Alexa 488-conjugated antibody, green) at NHBE cell surface (images merged in the bottom). Top: incubation protocol – CoV2-S_JN.1_-A647 1 h at 4°C for cell-surface attachment. **C,** PIX, but not ACE2 antibody, inhibits CoV2-S_JN.1_ attachment in human primary small airway epithelial cells in air-liquid interface (ALI) format. **Images:** sampled confocal images of Hoechst (gray, labeling cell nucleus), antibody-labeled Nucleocapsid protein (N Protein of CoV2-S_JN.1_, green), and Spike protein (anti-S2P6 of CoV2-S_JN.1_, red) in ALI cells were incubated with CoV2-S_JN.1_ for 1 h at 4°C (for viral cell-surface attachment, protocol shown at the top). Before virus incubation, cells were pretreated with 1) control solution (Ctrl), 2) Pixantrone (PIX), or 3) anti-ACE2 antibody (ACE2 Ab). Images were projection images. **Bar graphs:** quantification of the fluorescence of immunolabelled nucleocapsid protein (F_N protein_, upper) and spike (F_Spike_, lower) for ALI cells incubated with CoV2-S_JN.1_ for 1 h at 4°C in three pre-treated conditions stated above (Ctrl: 2 independent experiments; PIX: 2 independent experiments; ACE2 Ab: 2 independent experiments). ***: p < 0.001 (ANOVA test). **D,** Both PIX and ACE2 antibody inhibit CoV2-S_JN.1_ infection of ALI cells. **Images:** Immunolabelled N-protein images from ALI cells being incubated with viruses with a protocol for viral genome expression in three conditions, including 1) control (Ctrl), 2) pretreated with PIX (before virus incubation), and 3) pretreated with ACE2 Ab. The images are from the entire ALI culture using 20x objective (scale bar: 1 mm). Incubation protocol (shown at the top): CoV2-S_JN.1_ incubation for 1 h at 4°C, followed by incubation at 37°C for 24 h. **Bar graph:** quantification of the fluorescence of immunolabelled N protein (anti-Nucleocapsid protein antibody) for ALI cells being incubated with CoV2-S_JN.1_ with a protocol for viral genome expression (protocol shown in the left) in three conditions stated above (Ctrl: 3 repeats; PIX: 3 repeats; ACE2 Ab: 3 repeats). ***: p < 0.001 (ANOVA test).

Next, we validated our findings in a physiologically relevant human model with primary human airway epithelial cells cultured at air-liquid interface (ALI) format. Similar to results observed in NHBE cells (Fig. 7A, B), immunostaining for nucleocapsid protein and spike confirmed that PIX, but not anti-hACE2 antibody, substantially reduced surface-associated viral signals after the incubation protocol (4°C for 1 h) for virus cell-surface attachment (Figs. 7C, S10). Cell outlines were not clear at ALI format, because cells are densely packed in many layers (Fig. S10), making imaging difficult to resolve each cell’s outline and thus the virus distribution along the cell plasma membrane (Fig. 7C). However, this result reflects virus cell-surface attachment, because 1) the 4°C-for-1-h incubation protocol allowed for largely virus attachment at the cell surface as shown in Figs. 3 and 6, 2) inhibition of cell-surface HS-binding (Fig. 6), but not hACE2, substantially reduced the virus signal after the incubation for surface attachment (Fig. 7C), and 3) inhibition of either HS-binding or hACE2 substantially reduced the virus signal after an incubation protocol for virus infection (Fig. 7D).

For Fig. 7D, we assessed CoV2-S_JN.1_ infection in primary human small airway epithelial cells cultured at ALI format, which were incubated with CoV2-S_JN.1_ for 1 h at 4°C, followed by incubation at 37°C for 24 h for viral genome expression. Immunostaining for nucleocapsid protein revealed robust JN.1 infection in control cultures, whereas pre-treatment of PIX or anti-hACE2 antibody significantly reduced N protein signal (Fig. 7D). This result (Fig. 7D), together with the attachment data (Figs. 6, 7A-C), suggests that HS mediates initial cell-surface binding with CoV2-S_JN.1_, while hACE2 contributes at a downstream step required for infection.

## Discussion

### A two-receptor entry model – HS for attachment/endocytosis and ACE2 for downstream

We used a variety of advanced microscopic methods, including confocal, STED, MINFLUX, and EM imaging of pseudotyped (Figs. 1–3) and authentic SARS-CoV-2 (Figs. 4 and 5), HS, hACE2, as well as SARS-CoV-2-containing endocytic vesicles. This revealed HS as the SARS-CoV-2 receptor underlying virus cell-surface attachment and subsequent endocytic internalization, whereas hACE2, the generally believed SARS-CoV-2 receptor (*4*), is only required after virus attachment and endocytic internalization to promote viral genome expression (summarized in Fig. 5I). Clusters of ∼6-137 HS molecules, not single HS molecules, are the docking site for SARS-CoV-2 attachment (Figs. 3 and 5). The cell surface is covered by extremely dense clusters (∼1 cluster per 6 μm^2^) protruding from the plasma membrane by up to ∼410 nm (Fig. 5). This nanoscale arrangement – the tall, anionic HS-concentrated clusters densely covering the cell surface – may significantly enhance the chance that HS molecules in clusters bind multivalently to basic virion surface amino acids. We propose that HS outcompetes ACE2, which extends less than 10 nm from the plasma membrane (*40*), for virion attachment (*9, 21, 41, 42*).

Although hACE2 is not required for SARS-CoV-2 attachment and endocytosis, it is required for viral genome expression (Fig 1). It has been suggested that hACE2 binding changes S conformation to enable the protease cleavage of S needed for viral fusion (*4*). Cell-surface hACE2 co-internalized with SARS-CoV-2 in endosomes (Fig. 3I) may thus promote infection by changing S conformation, allowing endosomal cathepsin L to cleave S (Fig. 5I). Taken together, our results suggest a SARS-CoV-2 entry model with two receptors: HS clusters serve as the docking site for virus attachment and subsequent endocytic internalization, while hACE2 promotes downstream genome expression (Fig. 5I). We verified this model for authentic SARS-CoV-2 JN.1 subvariant binding on the cell-surface with HS and subsequent infection in primary human airway cells (Figs. 6, 7). Such cell surface attachment and infection were inhibited by a clinically used HS-binding agent, pixantrone (Fig. 7), suggesting that interfering with HS-binding is a key therapeutic strategy for COVID-19.

Our data suggest that, under physiological conditions in which HS is present at the cell surface, hACE2 is not essential for viral cell-surface attachment or endocytosis. We do not know whether hACE2 expression alone, in the absence of HS, can support viral cell-surface attachment and endocytosis. Although our data suggest that hACE2 functions downstream of endocytosis to facilitate viral fusion at the endosome for genome delivery to the cytosol, we do not know whether hACE2 binding with the virus occurs at the cell surface or endosomes. The binding may occur in both places, but not essential for virus attachment and endocytosis. Consistent with this possibility, HS may promote spike-dependent ACE2 super-cluster assembly at the cell surface and enhance SARS-CoV-2–associated syncytium formation, suggesting that HS may organize ACE2 nanoscale architecture in a cell–cell fusion context (*43*).

Our model (Fig. 5I) differs in three aspects from the generally accepted current model (*4, 9, 10*). First, the current model considers hACE2 essential in mediating viral attachment and initiation of endocytosis, whereas our model does not. Second, the current model considers HS as an attachment factor augmenting hACE2’s role in binding viruses for attachment and initiation of subsequent endocytosis, whereas our model considers HS as the receptor for viral attachment and subsequent endocytosis. Third, the current model considers virus-hACE2 binding and its facilitation by HS at the cell surface before endocytosis, whereas our model moves these actions downstream of endocytosis, likely in the endosomal membrane, the principal site of virion entry.

Although our SARS-CoV-2 entry model (Fig. 5I) differs from the widely accepted model, our findings do not necessarily contradict previous findings underpinning the dominant model. hACE2 binding with S in vitro (*5–7*) and its pivotal role in infection (*8*) led to the current view that hACE2 is the receptor for SARS-CoV-2 cell-surface attachment and subsequent entry into cells (*4*), despite the lack of direct observation of SARS-CoV-2 binding with hACE2 at the cell surfaces. Although light microscopic imaging showed S and hACE2 at the plasma membrane (*22, 44*), limited resolution precluded demonstrating a direct association. We clearly showed the cell-surface association of VSV-S and SARS-CoV-2 with HS, but not ACE2 (Figs. 3 and 5). This finding is consistent with previous studies showing HS binding with S *in vitro* and inhibition of SARS-CoV-2 infection by perturbing this binding with existing drugs or heparins, despite these results leading to proposing HS as an attachment factor facilitating S binding to the cell-surface ACE2 (*4, 9–11*). While some studies using genome-wide CRISPR screening to identify genes involved in SARS-CoV-2 reveal genes for HS biosynthesis, others do not (*45–50*). The negative result, which might reflect screen sensitivity, incomplete knockout, pathway redundancy, or viral dose/stringency, needs to be verified with specific gene knockout.

Our observation of the large size (115 - 789 nm) of vesicles containing 1-5 virions suggests that the endocytosis involved may form large vesicles. This form of endocytosis could be macropinocytosis, which can form large vesicles to engulf multiple virions. Consistent with this suggestion, a recent study showed that a macropinocytosis inhibitor, EIPA, reduces SARS-CoV-2 replication in Vero cells (*27*). Our finding that viral genome (*V-EGFP*) expression was largely inhibited (by ∼80%) or nearly abolished by inhibition of endocytosis or endosomal cathepsin L crucial for SARS-CoV-2 endosomal fusion (Fig. 2) suggests endocytic internalization as the major pathway for SARS-CoV-2 genome expression. While we cannot completely rule out a minor role for viral fusion at the plasma membrane in viral genome expression, our data indicate endocytosis following HS-mediated viral cell-surface attachment as the primary pathway for SARS-CoV-2 infection in the cells we studied. For other cells not examined in the present work, if TMPRSS2 is highly expressed, we could not rule out the possibility that the fusion pathway could also be dominant.

### The two-receptor model’s potential clinical relevance and broader significance

Interfering with HS binding has been suggested as a therapeutic strategy to prevent and treat many viral infections that depend on HS for entry, including COVID-19 (*1, 2, 9, 12*). Supporting this strategy, disrupting Spike–HS interactions, including inhibition by heparin and related glycans, reduces SARS-CoV-2 attachment/entry (*51*). Clinical evaluation of inhaled/nebulized unfractionated heparin has reported improved clinical outcomes without major bleeding signals, supporting the feasibility of targeting airway-surface HS interactions (*52*). HS mimetics, such as pixatimod (PG545), inhibit SARS-CoV-2 infection and exhibit greater potency than heparin in assays measuring inhibition of Spike/ACE2 engagement and viral infectivity (*53*). These reports support the translational potential of therapeutically interfering with virion–HS binding. However, this strategy has not been the focus for developing methods to prevent and treat COVID-19, likely because HS is considered only a regulator that is not essential for SARS-COV-2 entry. Our finding that HS is the attachment receptor re-emphasizes the importance of perturbing virus-HS binding, the first step of the viral entry, to efficiently block SARS-CoV-2 infection. Further supporting this view, inhibition of HS binding with a clinically used HS-binding agent, pixantrone, inhibits authentic SARS-CoV-2 JN.1 subvariant binding with HS on the cell-surface and infection in primary human airway cells (Figs. 6, 7). These results suggest a combinatorial anti-SARS-CoV-2 strategy: early HS blockade to prevent attachment combined with ACE2 targeting to inhibit post-attachment steps.

Analogous to SARS-CoV-2 (*4, 9–11*), without direct supporting evidence, HS has been considered a cell-surface attachment factor regulating other coronaviruses binding with their receptor hACE2 for attachment and subsequent cell entry (*2, 3, 12, 13*). This includes SARS-CoV, responsible for the SARS pandemic two decades ago, and human common cold coronavirus NL63 (*2, 3, 12, 13*). Our findings suggest that HS may also serve as the receptor for attachment and subsequent endocytosis of these CoVs. Perturbing virus-HS binding may be a broad anti-human CoV strategy.

Besides coronaviruses, HS is considered an attachment factor for and can bind to, hepatitis C virus, HIV, Ebola virus, human papillomavirus, and herpesviruses (*1*). For Ebola, HS has been proposed to deliver virions into endosomes (*54*). Our results implicate that HS may be a common cell-surface attachment receptor rather than a regulator for these viruses. This is because the anionic HS-concentrated clusters are densely packed at the cell surface and may extend hundreds of nanometers above the plasma membrane (Fig. 5) for easy access and strong binding with basic residues of approaching viruses’ surface proteins. Docking at the HS cluster may thus be a general strategy for many viruses to attach to the cell surface. Interfering with HS binding (*2, 4, 9, 12*) may constitute a general anti-viral strategy for not only HS/hACE2-dependent coronaviruses, but also many other HS-dependent viruses mentioned above. For SARS-CoV-2 and other HS-binding respiratory viruses, nasal spray delivery of HS competitors or heparinase may be an inexpensive but effective general anti-viral strategy.

## Acknowledgements

We thank Susan Cheng and Sandra Lara Moreira for EM technical support. This work was supported by NINDS Research Program (ZIA NS003009-15 and ZIA NS003105-10 to L.G.W.) and the Intramural Research Program of the National Institute of Allergy and Infectious Diseases National Institutes of Health (J.W.Y.) and the NIBIB (to A.J.J.), National Institutes of Health. The contributions of the NIH author(s) were made as part of their official duties as NIH federal employees, are in compliance with agency policy requirements, and are considered Works of the United States Government. However, the findings and conclusions presented in this paper are those of the author(s) and do not necessarily reflect the views of the NIH or the U.S. Department of Health and Human Services.

## Author contributions

S.H. X.W. and T.L. performed most experiments and participated in designing experiments; A.M., C.Y.C., S.H., J.M., and C.A.W. performed MINFLUX imaging and analysis. T.L., A.D.L. and I.K. and J.W.Y. performed experiments related to authentic SARS-CoV-2 and participated in designing experiments. R.S., Z.W. and A.J.J assisted S.H. for experiments. L.G.W. designed experiments, supervised the project, and wrote the manuscript with help from J.W.Y. and all other authors.

## Data and materials availability

All data needed to evaluate the conclusions in the paper are present in the paper and/or the Supplementary Information.

## Methods

### Cell lines, primary cells and air-liquid interface (ALI) airway epithelial cell culture

We used baby hamster kidney fibroblast BHK21 (BHK, ATCC, #CCL-10), human embryonic kidney 293T (293T, ThermoFisher Scientific, R70007), African green monkey kidney Vero E6 (Vero, ATCC, #C1008 [clone E6]), Hela (ATCC, #CCL-2), and human lung fibroblast MRC5 (ATCC, #CCL-171) cells. We generated BHK21 cells stably expressing human ACE2 (BHK_hACE2_) and 293T cells stably expressing human ACE2 (293T_hACE2_) using the Sleeping Beauty transposon plasmid expression system (*55*). All cell lines were cultured in Dulbecco’s modified Eagle’s medium (DMEM, Gibco, 11885084) supplemented with 10% fetal bovine serum (FBS, Gibco, 10082147) and incubated at 37°C and 5% CO_2_ in a humidified incubator. Wild-type CHO-K1 (ATCC, #CCL-61) and glycosaminoglycan (GAG)-deficient mutants CHO-pgsA-745 (ATCC, #CCL-2242) and CHO-pgsB-618 (ATCC, #CCL-2241) were maintained in F-12K medium (Gibco, 21127022) supplemented with 10% fetal bovine serum and 1% MEM NEAA (Gibco, 11140050) and incubated at 37°C and 5% CO_2_ in a humidified incubator.

We generated 293T cells stably expressing T7 polymerase and VSVG (293T-T7-VSVG) for vesicular stomatitis virus (VSV) virion production and Vero cells stably expressing human transmembrane serine protease 2 (Vero-TMPRSS2) for authentic SARS-CoV-2 virion production, using the Sleeping Beauty transposon plasmid expression system (*55*). Cell lines were confirmed to be mycoplasma-free using MycoStrip detection kit (InvivoGen, rep-mys-50).

Normal human bronchial/tracheal epithelial cells (Lonza, CC-2540) were expanded in PneumaCult-Ex Plus Basal medium (Stemcell, #05041) in flasks and incubated at 37°C and 5% CO_2_ in a humidified incubator. Differentiated primary human airway epithelial cultures were purchased from Epithelix (MucilAir) and maintained per the manufacturer’s instructions. Upon receipt, inserts were equilibrated for 24-48 h at 37 °C, 5% CO₂ with MucilAir™ medium (MA medium, Epithelix) in the basolateral chamber (typical volume 500–600 µL per insert), with no liquid on the apical side. MA medium was replaced 2–3 times per week. Before assays, the apical surface was gently rinsed twice with warm DPBS to remove accumulated mucus. Only well-differentiated tissues with robust ciliary beating and intact morphology were used.

### Generation of pseudo-typed SARS-CoV-2 virions: VSV*_EGFP_*-S-A647, VSV*_EGFP_*-S-A490, VSV*_EGFP_*-S_Omi_-A647 and Lenti*_EGFP_*-S_Omi_-A647

Generation of recombinant vesicular stomatitis virus (containing *EGFP*-coding RNA in the genome) pseudo-typed with SARS-CoV-2’s wild-type S-protein (S) of Wuhan-hu-1 strain with two modifications (VSV*_EGFP_*-S), the C-terminal 21 amino acids deletion and a furin cleavage site mutation R685S, was performed as previously described (*56*). Its use in tissue culture at biosafety level 2 was approved by the Institutional Biosafety Committee (IBC) at The National Institute of Allergy and Infectious Diseases (NIAID). Plasmid-based rescue of VSV*_EGFP_*-S was carried out as described previously with some modifications. Briefly, 293T-T7-VSVG cells in a 6-well plate were transfected with the VSV*_EGFP_*-S antigenome plasmid, along with plasmids expressing codon-optimized T7 polymerase (T7opt in pCAGGS, addgene, 65974), and VSV N, P, L and G-expressing plasmids (Addgene, 64087, 64088, 64085 and 8454), using TransIT-LT1 transfection reagent. The medium was exchanged with fresh DMEM containing doxycycline (1 μg/mL, for VSVG expression). EGFP expression was monitored every day for 2-3 days. Transfected cells with EGFP-positive clusters and typical cytopathic effect (CPE) were collected as passage 0 and then inoculated to fresh BHK_hACE2_ cells to propagate as viral stock P1, which underwent further propagations to generate the VSV*_EGFP_*-S virions in Vero or BHK_hACE2_ cells. Viral passages were subjected to Sanger and next-generation sequencing to verify the S gene or the viral genome.

VSV*_EGFP_*-S virus was amplified in Vero or BHK_hACE2_ cells. The amplified VSV*_EGFP_*-S virus was either concentrated with an ultrafast centrifuge, or Lenti-X concentrator (Takara, #631232) according to the manufacturer’s protocol. The protein level of VSV*_EGFP_*-S virus was ∼6 μg/μl, which was measured with a NanoDrop spectrophotometer (ND-1000).

Purified VSV*_EGFP_*-S was incubated with Alexa Fluor 647 NHS ester (A647, 7.7 μM, Invitrogen, #A20006) with gentle agitation for ∼1.5 h at 22-24°C, which led to A647 conjugation with primary amines of superficial envelope proteins of virus (VSV*_EGFP_*-S-A647). Free A647 dyes were removed with 7 k molecular weight cutoff desalting columns (7 k MWCO, Thermo, #89882). Viruses were aliquoted and stored at -80°C for further experiments. The resulting virion, VSV*_EGFP_*-S-A647, could be observed with fluorescent imaging of A647 (Fig. 1A).

In some experiments, we replaced wild-type S protein in VSV*_EGFP_*-S-A647 with the Omicron variant to produce the VSV*_EGFP_*-S_Omi_-A647 virion; we replaced A647 in VSV*_EGFP_*-S-A647 with Atto 490LS NHS ester (A490, 63 μM, Atto-Tec GmbH, #AD 490LS-25) to produce the VSV*_EGFP_*-S-A490 virion.

Lentivirus (with *EGFP*-coding sequence in the genome) pseudo-typed with S protein Omicron variant (Lenti*_EGFP_*-S_Omi_) was purchased from BPS Bioscience (#78624-2). Lenti*_EGFP_*-S_Omi_ was incubated with A647 (7.7 μM) for ∼1.5 h at 22-24°C, and purified with desalting columns, which led to A647 conjugation with primary amines of superficial envelope proteins. The resulting virion was termed Lenti*_EGFP_*-S_Omi_-A647.

### Generation of recombinant SARS-CoV-2 Omicron variant

The generation of recombinant SARS-CoV-2 Omicron variant (CoV2-S_Omi_) and its use in tissue culture at biosafety level 3 were approved by the IBC and the Dual Use Research of Concern Institutional Review Entity (DURC-IRE) at NIAID. Genetic modification inserted a mCherry gene to the nucleoprotein via a 2A linker, derived from a bacterial artificial chromosome-based SARS-CoV-2 reverse genetic system (referred to as CoV2*_mCh_*-S_omi_) (*56*). Virus rescue experiments were performed as previously described (*56*). Briefly, BHK_ACE2_ cells were transfected with pBAC-SARS-CoV-2 and media were changed with DMEM containing 2% FBS 6 h later. mCherry-positive cells were detached and collected along with supernatant during 48-72 h and stored at - 80°C as the first stock. The stock was centrifuged to remove cell debris and used to infect fresh Vero-TMPRSS2 cells for 48-72 h. The supernatant was collected and stored at -80°C as the second stock. After the rescued virus in the second stock was confirmed by Sanger sequencing, the virus in the first stock was allowed to undergo two rounds of propagation to generate the CoV2*_mCh_*-S_Omi_ virions. The viral genome sequence was verified by next-generation sequencing before virus labeling with fluorescent dyes.

Purified CoV2*_mCh_*-S_omi_ was incubated with Alexa Fluor 647 NHS ester (A647, 7.7 μM, Invitrogen, #A20006) for ∼1.5 h at 22-24°C, which led to A647 conjugation with primary amines of superficial envelope proteins. The resulting virion, CoV2*_mCh_*-S_omi_-A647, was purified with desalting columns (7 k MWCO, Thermo, #89882). CoV2*_mCh_*-S_omi_-A647 could be observed with confocal, STED, and MINFLUX imaging of A647 (Figs. 4 and 5). In some experiments, we replaced A647 in CoV2*_mCh_*-S_omi_-A647 with Atto 490LS NHS ester (A490, Atto-Tec GmbH, #AD 490LS) to produce CoV2*_mCh_*-S_omi_-A490.

### Preparation of SARS-CoV-2 JN.1 variant and fluorescent labeling

A clinically isolated JN.1 subvariant (BA.2.86.1.1) of SARS-CoV-2 (CoV2-S_JN.1_) was propagated under BSL-3 conditions. Purified CoV2-S_JN.1_ was incubated with Alexa Fluor 647 NHS ester (A647, 7.7 μM, Invitrogen, #A20006) for ∼1.5 h at 22-24°C, which led to A647 conjugation with primary amines of superficial envelope proteins. The resulting virion, CoV2-S_JN.1_-A647, was purified with desalting columns (7 k MWCO, Thermo, #89882). For some binding and infection assays, unlabeled JN.1 stocks were used.

### Plasmids, transfection, cell plating, and fluorescent dyes

The hACE2-TagBFP plasmid was purchased from Addgene (#164219). The mutant Dynamin 2-K44A-mCherry plasmid was created from Dynamin 2-mCherry (Addgene, #27689). The PH_mNG_ (phospholipase C delta PH domain attached with mNeonGreen) construct was created by replacing the EGFP tag of PH-EGFP (obtained from Dr. Tamas Balla) with mNeonGreen (Allele Biotechnology) (*57*).

Cells were transfected with Lipofectamine LTX reagent (ThermoFisher Scientific, #A12621) according to the manufacturer’s protocol and plated on the collagen-coated glass bottom of dishes (MatTek, #P35GCol-1.5-14-C). Images were taken 48 hours after transfection.

We incubated cells with 50 μM A490 for 1 h to identify internalized vesicles with confocal imaging. We incubated cells with 500-750 μM A490 for 3 min for the measurement of the internalized vesicle size with STED microscopy.

### Immunostaining for confocal or STED imaging

Cells were fixed with 4% paraformaldehyde, blocked with 2% BSA in PBS solution for 1 h at 22-24°C, and subsequently incubated with primary and secondary antibodies. Primary antibodies were diluted in PBS containing 2% BSA and incubated with cells at 4°C for 16-20 h. After 3 times wash in PBS, cells were incubated with fluorescence-conjugated secondary antibodies for 1 h at 22-24°C. After washing 3 times with PBS, cells were mounted on confocal and STED microscopes to acquire images. Primary antibodies used here included goat anti-ACE2 (1:500, R&D, #AF933), mouse anti-heparan sulfate (1:500, Amsbio, 10E4), human anti-S2P6 (1:500) and rabbit anti-nucleocapsid (1:500, GeneTex, GTX135357); secondary antibodies included Alexa Fluor 647-labeled donkey anti-goat IgG (Thermo, #A21447) and Alexa Fluor 568-labeled goat anti-human IgG (Thermo, #A21090), Alexa Fluor 488-labeled goat anti-mouse IgG (Thermo, #A11029), Alexa Fluor 488-labeled goat anti-rabbit IgG (Thermo, #A11028), Alexa Fluor 488-labeled donkey anti-mouse IgM (Jackson, #715-545-140).

### Confocal imaging

Confocal images were acquired with Leica TCS SP8 inverted confocal microscope (Leica, Germany) equipped with a × 63/1.40 oil immersion objective. Alexa 647 (A647), EGFP, A490 and mCherry were excited by solid-state lasers with wavelength of 638 nm (30 mW output), 488 nm (20 mW output), 514 nm (20 mW output) and 552 nm (20 mW output) respectively, and their fluorescence signals were collected with Leica HyD detectors at 650-795 nm, 500-550 nm, 560-630 nm, and 650-795 nm, respectively. To visualize the viruses labeled by A647 or A490, 638 nm or 514 nm laser was set at 0.2–2% or 5% of the maximum power; to detect virus *EGFP* or *mCherry* expression, 488 nm or 552 nm excitation laser was set at 0.5–2% or 50% of the maximum power, respectively.

To measure virus attachment or endocytosis, or virus genome (*EGFP/mCherry*) expression, XY-plane scanning along the Z-axis every 1 μm was performed (XY/Z_stack_ scanning), and the projected fluorescence were quantified with Fiji software for each cell to quantify all the particles at cell surface or inside the cell, or the total fluorescence.

To measure HS puncta number at cell surface or total area of cell surface, XY/Z_stack_ scanning with a Z-axis interval of 0.5-1 µm was performed to obtain the images of the entire cell. 3D reconstruction was generated and analyzed using Spots or Surfaces function with Imaris software (Oxford Instruments).

### STED imaging

The STED images were acquired with XY-plane scanning mode at a fixed Z-axis focal plane. STED images were acquired with the Abberior Expertline system based on an inverted Olympus IX83 microscope that is equipped with a UPlanSApo 100 x 1.4 NA oil immersion objective. To measure virus (VSV*_EGFP_*-S-A647 or CoV2*_mCherry_*-S_Omi_-A647) endocytosis with bath A490 (Fig. 2A-D and Fig. 4E), A647 and A490 were excited at 640 nm and 512 nm respectively, and both collected at 650-765 nm. STED depletion was conducted with 775 nm depletion beam at 5-10% of the maximum depletion power. XY/Z_stack_ imaging with Z-axis interval of 200 nm was performed.

To measure viral colocalization with heparan sulfate and hACE2 on the cell surface, cells were incubated with VSV*_EGFP_*-S-A490 at 4°C for 1 h, fixed with 4% paraformaldehyde, and stained with primary and secondary antibodies. For STED imaging of VSV*_EGFP_*-S-A490 and heparan sulfate (Fig. 4C), mouse anti-heparan sulfate (1:500, Amsbio, 10E4) and Alexa fluor 488-labeled donkey anti-mouse IgM (Jackson, #715-545-140) were used to label heparan sulfate. We first acquired VSV*_EGFP_*-S-A490 STED images with 775 nm depletion laser together with heparan sulfate-Alexa 488 (A488) confocal images, and then both STED and confocal images of heparan sulfate-Alexa 488 were acquired with 595 nm depletion laser. VSV*_EGFP_*-S-A490 and heparan sulfate-A488 were excited by 640 nm and 512 nm laser, respectively, and their fluorescence signals were collected at 650-765 nm and 528-585 nm, respectively. The power of 640-nm laser was set at 3-5% of the maximum (maximum power: 1 mW); the corresponding 775 nm depletion laser power was set at 5-10% of the maximum (maximum power: 3 W). The power of 512-nm laser was set at 3-5% of the maximum (maximum power: 1 mW); the corresponding 595 nm depletion laser power was set at 5-10% of the maximum (maximum power: 700 mW).

For imaging of VSV*_EGFP_*-S-A490 and hACE2 (Fig. 4H-I), goat anti-ACE2 (1:500, R&D, #AF933) and Alexa fluor 647-labeled donkey anti-goat IgG (Thermo, #A-21447) were used to label hACE2. We acquired VSV*_EGFP_*-S-A490 and ACE2-Alexa 647 (A647) STED images with 775 nm depletion laser. VSV*_EGFP_*-S-A490 and ACE2-A647 were sequentially excited by 512 nm and 640 nm laser, respectively, and their fluorescence signals were sequentially collected at 650-765 nm. The power of 512-nm and 640-nm laser were set at 3-5% of the maximum and the 775 nm depletion laser power was set at 5-10% of the maximum.

STED images were deconvolved using Huygens software (Scientific Volume Imaging) as described previously (*58*) and analyzed with Image J and LAS X (Leica).

### Electron microscopy

BHK_hACE2_ cells were incubated with VSV*_EGFP_*-S at 4°C for 1 h, then transferred to 37°C for 15 min (to track VSV*_EGFP_*-S in intracellular vesicles, Fig. 2E). BHK_hACE2_ cells were fixed with 4% glutaraldehyde (freshly prepared, Electron microscopy sciences, Hatfield, PA) in 0.1 N Na-cacodylate buffer for 1 h at 22-24°C, and stored in 4°C refrigerator till processing. Cells were washed with 0.1 N cacodylate buffer and treated with 1% OsO4 in cacodylate buffer at 4°C for 1 h, then with 0.25% uranyl acetate (pH 5.0) at 4°C for 16-20 h. Next, cells were dehydrated with ethanol, and embedded in epoxy resin. Thin (70 nm) sections were counterstained with uranyl acetate and lead citrate and then imaged in a JEOL 200 CX TEM. Images were captured with a CCD digital camera system (XR-100 from AMT, Danvers, MA) at a primary magnification of ×10,000-30,000.

### Reagents and their application procedures

Dynasore (DnS, 80 μM, Millipore Sigma) (*59, 60*) was applied to the cell media 30 min before the virus was applied, and was continuously present for another 1 h during which the virus was added to the cell media (see Fig. 2F, G for the viral incubation protocol). Cathepsin inhibitor 1 (CPS Inh, 10 μM, Selleckchem, S2847) (*31–34*) was applied to the cell media 30 min before the virus was applied; CPS Inh was continuously present for another 1 h (Fig. 2H, left) during which the virus was added to the cell media, or for another 24 h (Fig. 2H-I, right) during which the virus was added for 1 h and then washed out.

HPRase (*35*), the heparinase I/II/III mixture containing heparinase I (3.2 units, NEB, #P0735S), heparinase II (1 units, NEB, #P0736S) and heparinase III (0.19 units, NEB, #P0737S), was applied to cells in FBS-free DMEM at 30°C for 1 h before the virus was applied. After washing out, cells were either fixed for immunostaining of heparan sulfate (Fig. 3D) and hACE2 or incubated with viruses at 4°C for 1 h. Virus attachment was measured after 1 h incubation at 4°C (Fig. 3E and Fig. 4F). Cells were then put back to 37°C incubator for another 1 h to measure the virus uptake (Fig. 3F), or for another 24 h to measure the *EGFP* gene expression (Fig. 3G).

Pixantrone (PIX, 20 µM, Selleck), a HS-binding agent under clinical trial (*56*), or anti-human ACE2 monoclonal antibody (ACE2 Ab, 20 µg/mL, R&D, #AF933) was applied to the cells 30 min before the virus was applied and was continuously present for another 1 h during which the virus was added to the cell media (see Fig. 7A, C for the viral incubation protocol). For ALI tissues, excess apical mucus was gently removed with warm PBS (1–2 rinses) prior to incubation. Plates with inserts were pre-chilled and PIX or ACE2 Ab was applied to the apical surface and maintained during attachment or infection. After incubation, cells were washed with ice-cold medium to remove unbound virions. Inserts were immediately fixed with 4% PFA for immunostaining for binding assay (Fig. 7C) or changed to culture at 37 °C for 24 h, then fixed with 4% PFA for immunostaining for infection assay (Fig. 7D).

### Flow cytometry

To measure viral attachment, BHK_hACE2_ or BHK cells were incubated with or without VSV*_EGFP_*-S-A647 at 4°C for 1 h, washed three times with PBS, incubated with ice-cold DPBS (Gibco) supplemented with EDTA solution (5 mM, QualityBiologicals) for 30 min. Cells were rigorously pipetted to generate single-cell suspensions (Fig. S6). To measure viral infection on linked samples, BHK_hACE2_ or BHK cells were then incubated with or without VSV*_EGFP_*-S-A647 at 37°C for 24 h. After incubation, cells were washed with PBS three times and resuspended in DPBS (Gibco) supplemented with BSA (0.1%) (Fig. S2). Flow cytometry was performed on BD FACSCelesta Cell Analyzer with BD FACSDiva software version 9.0. Initial data processing was done with FlowJo version 10.10. For all flow cytometry experiments, we did not include trypsin for digestion to avoid the potential impact of trypsin in cleaving proteins attached to heparan sulfate (*61*).

### Verification of VSV*_EGFP_*-S-A647 in vitro

To visualize VSV*_EGFP_*-S (Fig. 1A), a glass-bottomed cell culture dish was coated with lectin (Sigma) at 37°C for 1 h, and then 10 µl VSV*_EGFP_*-S-A647 was spread onto glass and followed by fixation with 2% paraformaldehyde at 22-24°C for 30 min. After washing, 2% bovine serum albumin (BSA)/PBS was used for block and antibody staining. VSV*_EGFP_*-S-A647 were incubated with anti-spike antibody (1A9, 5 µg/ml, Cat. No. GTX632604, GeneTex) targeting S2 subunit at 4°C for 16-20 h, followed by wash and 1 h of incubation with Alexa Fluor 488-labeled donkey anti-mouse IgG (Abcam). After the final wash, images (e.g., Fig. 1A) were acquired under a Leica TCS SP8 inverted confocal microscope (Leica, Germany) equipped with a × 63/1.40 oil immersion objective.

### MINFLUX nanoscopy

#### MINFLUX data acquisition

MINFLUX imaging was performed on a commercial 3D MINFLUX microscope that was operated by the Imspector software with MINFLUX drivers (Abberior Instruments) (*15*). Generally, fields of view with multiple gold beads were chosen and locked in a 3D set position for active sample stabilization using the near-infrared scattering from gold beads and active feedback correction via the piezo stage. It was ensured that the standard deviation of the sample position relative to the stabilization set point was less than 2 nm in all directions during measurements. Fields of view were chosen close to the coverslip surface at the bottom of cells. Before starting the MINFLUX data acquisition, the fluorophores were driven into the dark state (cyanine-thiol adduct) using iterative confocal scans with the 640 nm excitation laser and a power between 8-15% (maximum power at periscope: 1.94 mW). The sample was imaged with the standard MINFLUX imaging sequence provided by the manufacturer using 10% fixed laser power. During the MINFLUX measurement, the 405 nm activation laser power was ramped up slowly from 0% to 50% over several hours (maximum power at periscope: 27 μW). Samples were usually imaged for ∼2-6 hours.

#### MINFLUX data analysis

The raw final valid molecule position estimates were exported directly from the MINFLUX Imspector interface as a .mat file. Custom MATLAB analysis software was then used to identify and segregate clusters of localizations. The data were filtered to remove trace IDs (TIDs, group of localizations originating from the same fluorophore emission burst) with a standard deviation of more than 10 nm and less than three localizations per trace (*62, 63*).

For the estimate of the copy number of HS labelled with HS-antibody-A647, the TIDs per purified HS molecule *in vitro* were counted and used to normalize the number of TIDs per HS cluster at the cell-surface, assuming an unchanged A647 emission behavior between measurements *in vitro* and on the cell surface. Here, we did not count the molecules directly using the number of TIDs per cell-surface HS cluster, due to the reversibility of A647 photoswitching which would lead to overcounting. The diameter of each cluster of localizations was estimated by averaging the height and the width of an ellipse fit [(height + width)/2, Figs. S9A and S9D]. The diameter may refer to localizations for a single HS molecule in vitro, a cell-surface HS cluster, or a virion.

### Sample preparation for MINFLUX imaging

#### Preparing heparan sulfate on coverslips

For the attachment of heparan sulfate to coverslips, 400 µl heparan sulfate (Sigma-Aldrich, H7640) solution (1 µg/µl) was added to the glass-bottom dishes coated with collagen (MatTek, P35GCol-1.5-14-C) and allowed to stay at 4 °C for 16-20 h to adhere to the glass surface. After washing with PBS, the heparan sulfate on coverslips was blocked with 2% BSA in PBS for 1 h at 22-24 °C, followed by 1 h incubation of mouse anti-heparan sulfate (1:500, Amsbio, 10E4). Dishes with heparan sulfate were washed with PBS and incubated with a secondary goat anti-mouse IgG antibody conjugated with Flux 647 (1:500, Abberior) for 1 h. The heparan sulfate dishes were ready for the next step after washing three times with PBS.

Immediately prior to the confocal and MINFLUX imaging, the dish with heparan sulfate on the glass bottom was incubated with 200 µl undiluted gold bead solution (Gold nanoparticles 150 nm, BBI solutions, SKU: EM.GC150) for 5 min. After rinsing off the gold bead solution, the dish was washed with PBS for several times. An isotonic imaging buffer containing 200 mM Tris-HCl (pH 8.0), 10 mM NaCl, 10% (w/v) glucose, 63 µg/ml bovine liver catalase (Sigma-Aldrich, C1345), 393 µg/ml glucose oxidase from *Aspergillus niger* (type VII, Sigma-Aldrich, G2133) and 12.6 mM cysteamine (Sigma-Aldrich, 30070) was added to the glass bottom of the dish. A 25-mm coverslip (Neuvitro) was placed above the glass bottom to generate a reaction cavity, excessive imaging buffer was removed to seal the cavity before mounting the dish onto the microscope. The coverslips (above the glass bottom dishes) were sealed with Elite double 22 dental epoxy (Zhermack).

#### Preparing cells for MINFLUX imaging

BHK_hACE2_ cells were transfected with PH_mNG_ plasmid and plated on glass bottom dishes (MatTek, P35GCol-1.5-14-C). The transfected cells were allowed to express PH_mNG_ for 24-48 h. For MINFLUX imaging of heparan sulfate (Fig. 5B-D), cells were incubated with CoV2*_mCh_*-S_Omi_-A490 at 4 °C for 1 h (for confocal imaging), fixed with 4% paraformaldehyde, and then stained with mouse anti-heparan sulfate antibody and Flux 647-conjugated goat anti-mouse secondary antibody (HS-Ab-F647, for MINFLUX imaging). For MINFLUX imaging of CoV2*_mCh_*-S_Omi_-A647 (Fig. 5G-H), cells were incubated with CoV2*_mCh_*-S_Omi_-A647 at 4 °C for 1 h, and then fixed with 4% paraformaldehyde. Immediately prior to MINFLUX imaging, the dish was incubated with 200 µl undiluted gold bead solution (Gold nanoparticles 150 nm, BBI solutions, SKU: EM.GC150) for 5 min. After rinsing off the gold bead solution, the dish was washed with PBS for several times. An isotonic imaging buffer containing 200 mM Tris-HCl (pH 8.0), 10 mM NaCl, 10% (w/v) glucose, 63 µg/ml bovine liver catalase (Sigma-Aldrich, C1345), 393 µg/ml glucose oxidase from *Aspergillus niger* (type VII, Sigma-Aldrich, G2133) and 12.6 mM cysteamine (Sigma-Aldrich, 30070) was added to the glass bottom to generate a reaction cavity. The coverslips (above the glass bottom dishes) were sealed with Elite double 22 dental epoxy (Zhermack).

### Statistics

Data were presented as mean ± s.e.m. Replicates are indicated in results and figure legends. N represents the number of cells, A490-labeled vesicle spots, endosomes, viral spots, ACE2 or HS spots, HS clusters or experiments as indicated in results and figure legends. The statistical significance was tested with the unpaired Student’s t-test.

## Supplementary Information

**Figure S1.**
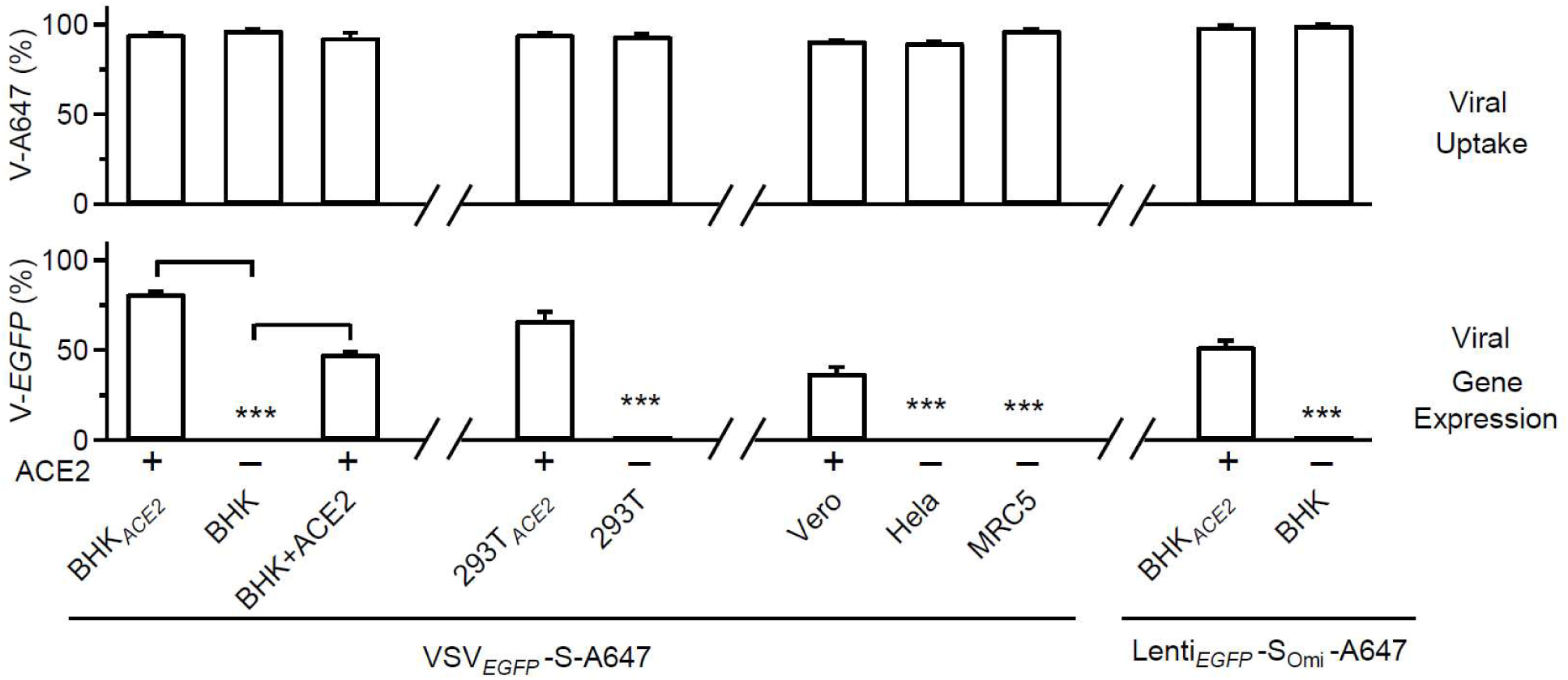
SARS-CoV-2 pseudovirus gene expression, but not internalization, depends on hACE2. The percentage (%) of cells containing VSV*_EGFP_*-S-A647’s V-A647 (%, mean ± s.e.m., upper) and *`V-EGFP`* (F*_EGFP_*, mean ± s.e.m., lower) in BHK_hACE2_ (3 experiments, 471 cells), BHK (3 experiments, 370 cells), BHK+hACE2 (3 experiments, 515 cells). 293T_hACE2_ 3 experiments, 404 cells), 293T (3 experiments, 234 cells), Vero (3 experiments, 295 cells), Hela (3 experiments, 305 cells), and MRC5 (3 experiments, 237 cells) cells. Lenti*_EGFP_*-S_Omi_-A647 virus’ V-A647 and *`V-EGFP`* in BHK_hACE2_ (3 experiments, 364 cells) and BHK (3 experiments, 335 cells) cells are also plotted. The virus incubation protocol is shown in Fig. 1B (1 h viral incubation, 24 h washout). The percentage of cells containing V-A647 or *`V-EGFP`* refers to cells identified in the bright field. ***: p < 0.001, t-test.

**Figure S2.**
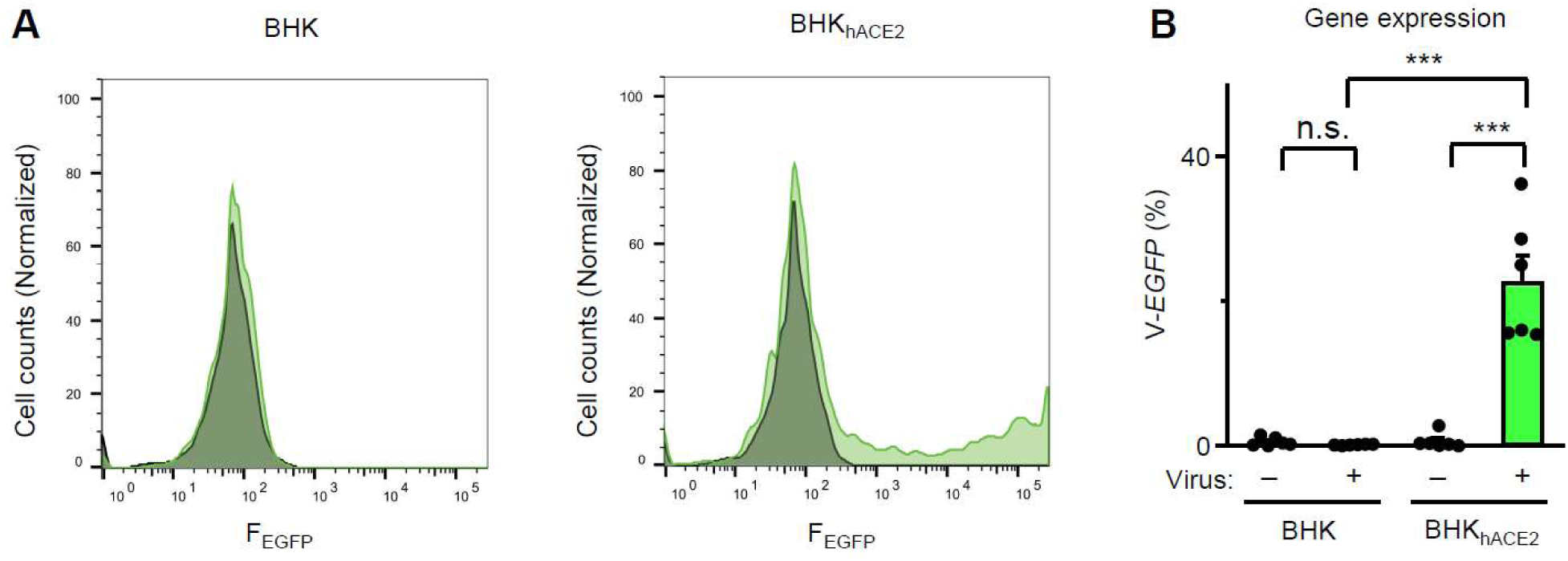
hACE2-dependent *V-EGFP* expression detected with flow cytometry. **A,** Cell counts (normalized to mode) plotted versus the EGFP fluorescence intensity (F_EGFP_) in BHK (left) and BHK_hACE2_ (right) cells incubated with VSV*_EGFP_*-S-A647 (green) or without VSV*_EGFP_*-S-A647 (black, control) at 37°C for 18 h. Ice-cold DPBS supplemented with EDTA solution was used instead of trypsinization, to avoid potential digestion of heparan sulfate proteoglycans. **B,** Percentage of cells expressing EGFP (*`V-EGFP`*) as detected by flow cytometry in BHK cells (biological triplicates for each experiment, 2 experiments) and in BHK_hACE2_ cells (biological triplicates for each experiment, 2 experiments) incubated with VSV*_EGFP_*-S-A647 (virus +) or without VSV*_EGFP_*-S-A647 (virus –, control) at 37°C for 18 h. ***: p < 0.001 (ANOVA test); n.s.: p > 0.05.

**Figure S3.**
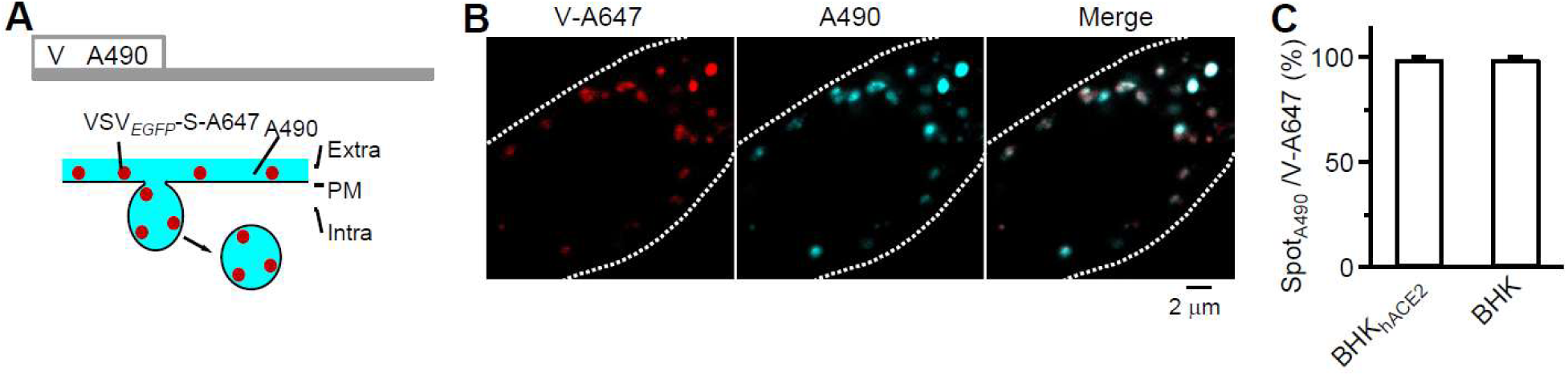
Uptake of both VSV*_EGFP_*-S-A647 and the bath A490 into the cytosol reflects viral endocytosis. **A,** Top: incubation protocol – VSV*_EGFP_*-S-A647 plus 50 μM A490 for 1 h at 37°C, followed by 24 h washout at 37°C. Bottom: strategy for labeling endocytic vesicles taking up PM-attached virions (VSV*_EGFP_*-S-A647, red) with a fluorescent dye in the bath (A490, cyan). **B,** Confocal images showing V-A647 spot (red dots) colocalization with A490 spots (green) in the cytosol. White dash line: cell outline from bright field images. **C,** The percentage of cytosolic V-A647 spots colocalized with A490 spots in BHK_hACE2_ (27 cells, 3 experiments) and BHK cells (21 cells, 3 experiments) after the incubation protocol shown in panel A.

**Figure S4.**
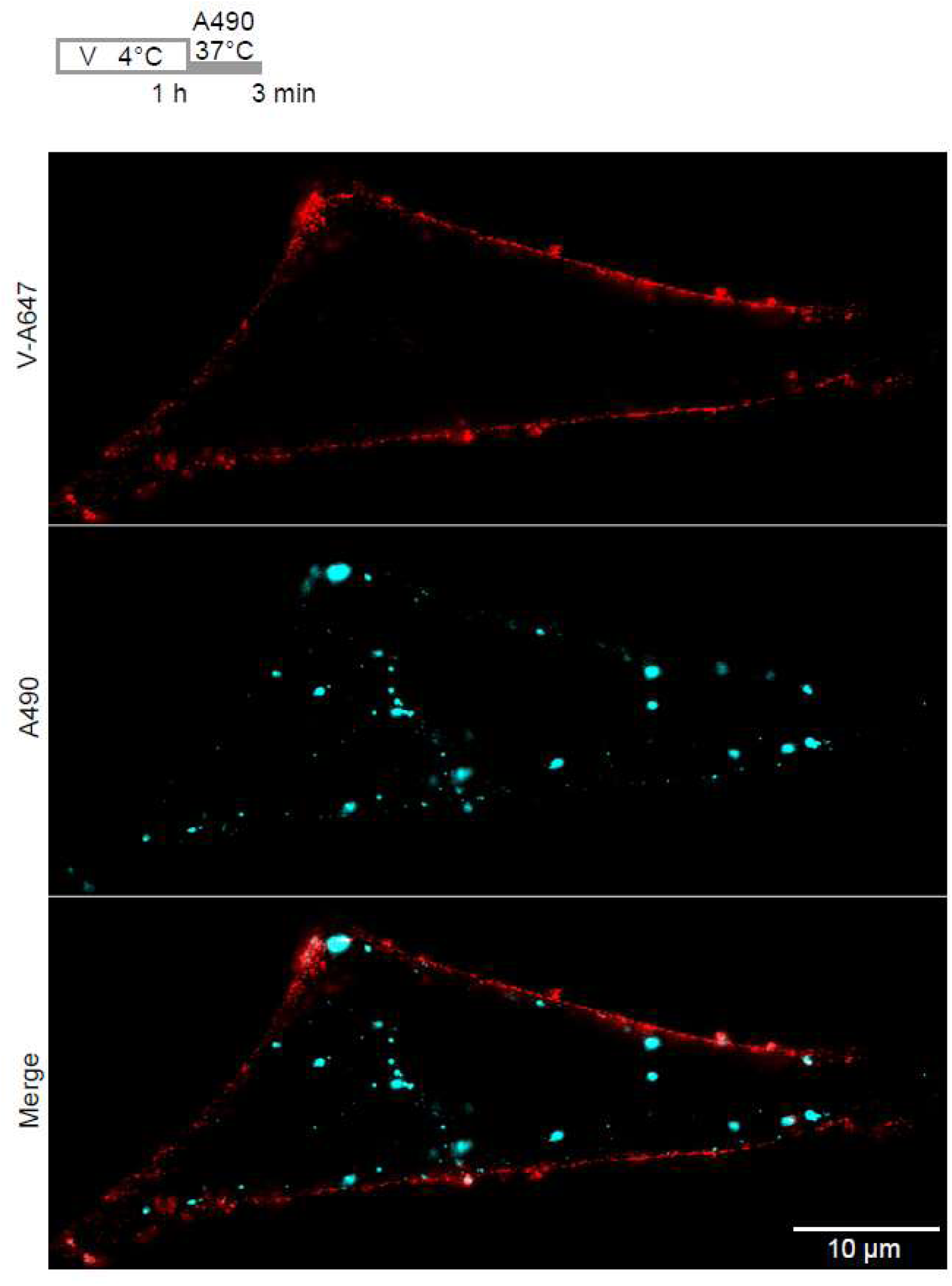
Virus cell-surface attachment labels the cell outline. STED images showing most V-A647 spots (red dots) decorating the outline of the cell plasma membrane and A490 spots (cyan) within the outline. Incubation protocol is shown on the top: 1 h VSV*_EGFP_*-S-A647 at 4°C, 3 min washout with 500 μM A490.

**Figure S5.**
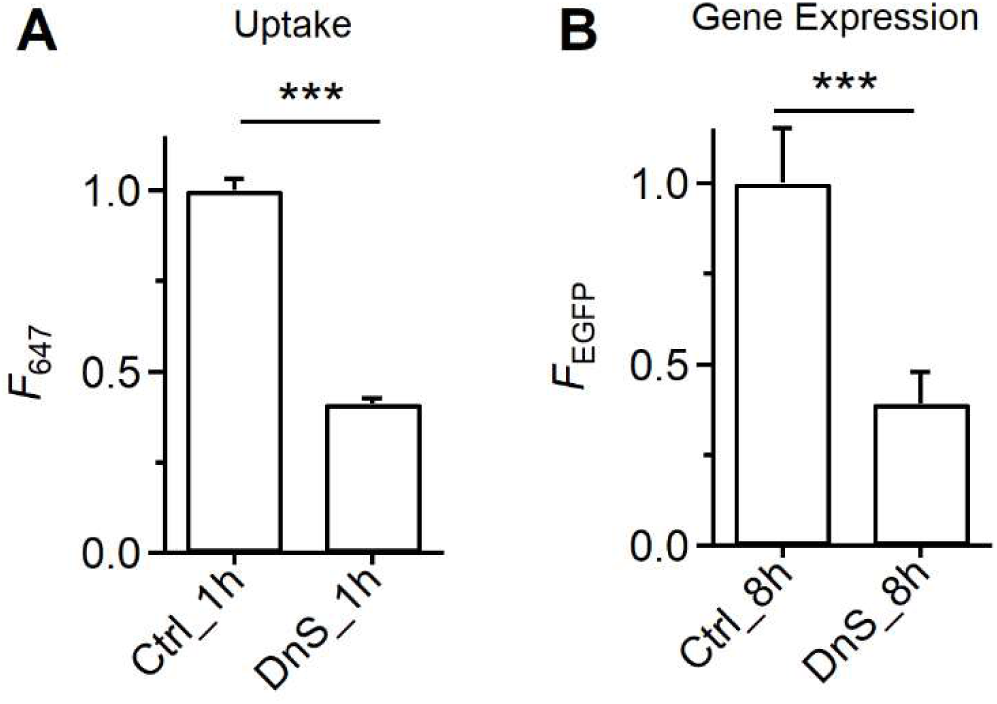
Dynasore reduces VSV*_EGFP_*-S viral uptake and gene expression. **A,** VSV*_EGFP_*-S-A647’s *`F`*_647_ in cells in control (Ctrl_1h, 256 cells) and treated with dynasore (DnS_1h, 248 cells) for 1 h, reflecting viral uptake. BHK_hACE2_ cells were treated with control (Ctrl) or dynasore (DnS, 80 μM, bath) with VSV*_EGFP_*-S-A647 at 37°C for 1 h for confocal images. Data are normalized to control group. ***: p < 0.001, t-test. **B,** VSV*_EGFP_*-S-A647’s *`F`_EGFP_* in cells in control (Ctrl_8h, 316 cells) and treated with dynasore (DnS_8h, 227 cells) for 8 h, reflecting viral gene expression. BHK_hACE2_ cells were treated with control (Ctrl) or dynasore (DnS, 80 μM, bath) with VSV*_EGFP_*-S-A647 at 37°C for 1 h, washout and imaged at 8 h. Data are normalized to control groups. ***: p < 0.001, t-test.

**Figure S6.**
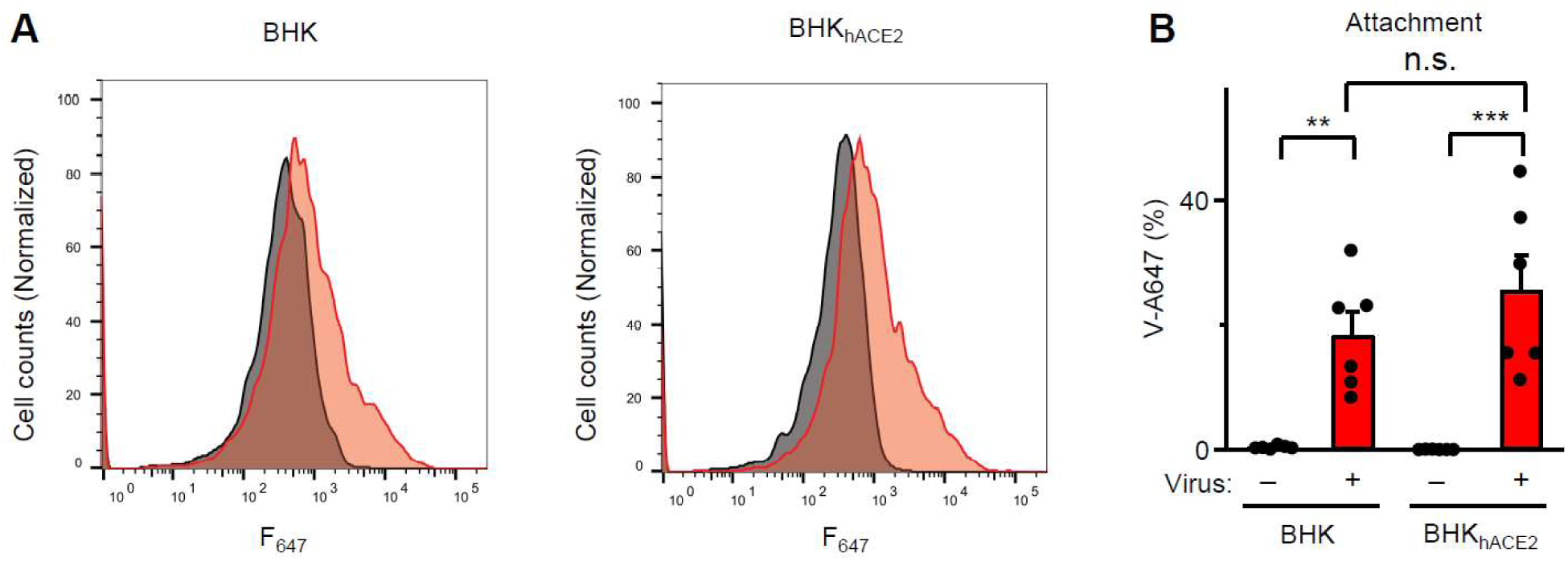
hACE2-independent viral attachment detected with flow cytometry. **A,** Cell counts (normalized to mode) plotted versus fluorescence intensity of VSV*_EGFP_*-S-A647’s V-A647 (F_647_) in BHK (left) and BHK_hACE2_ (right) cells incubated with VSV*_EGFP_*-S-A647 (red) or without VSV*_EGFP_*-S-A647 (black, control) at 4°C for 1 h. Ice-cold DPBS supplemented with EDTA solution was used instead of trypsinization, to avoid potential digestion of heparan sulfate proteoglycans. **B,** Percentage of cells associated with VSV*_EGFP_*-S-A647’ A647 (V-A647) detected with flow cytometry in BHK cells (biological triplicates for each experiment, 2 experiments) and in BHK_hACE2_ cells (biological triplicates for each experiment, 2 experiments) incubated with VSV*_EGFP_*-S-A647 (virus +) or without VSV*_EGFP_*-S-A647 (virus –, control) at 4°C for 1 h. **: p < 0.01; ***: p < 0.001 (ANOVA test); n.s.: p > 0.05.

**Figure S7.**
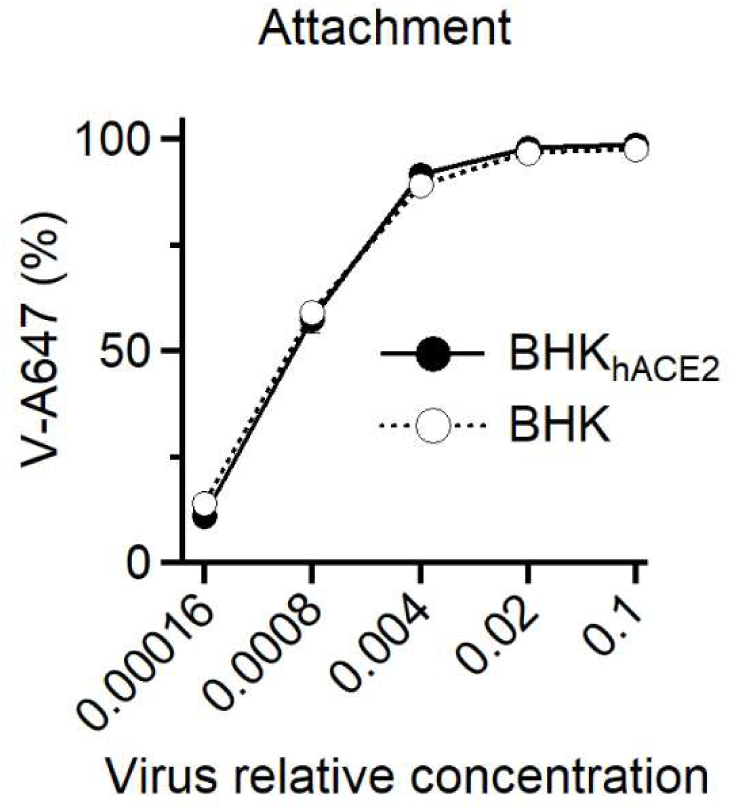
hACE2-independent viral attachment detected across a virus dilution series. Percentage of cells associated with VSV*_EGFP_*-S-A647’ A647 (V-A647) detected with flow cytometry in BHK and in BHK_hACE2_ cells incubated with VSV*_EGFP_*-S-A647 at 4°C for 1 h across different virus concentrations generated by serial dilution. Similar attachment was observed in BHK and BHK_hACE2_ cells across the tested concentration range, indicating that VSV-S cell-surface attachment is largely independent of hACE2.

**Figure S8.**
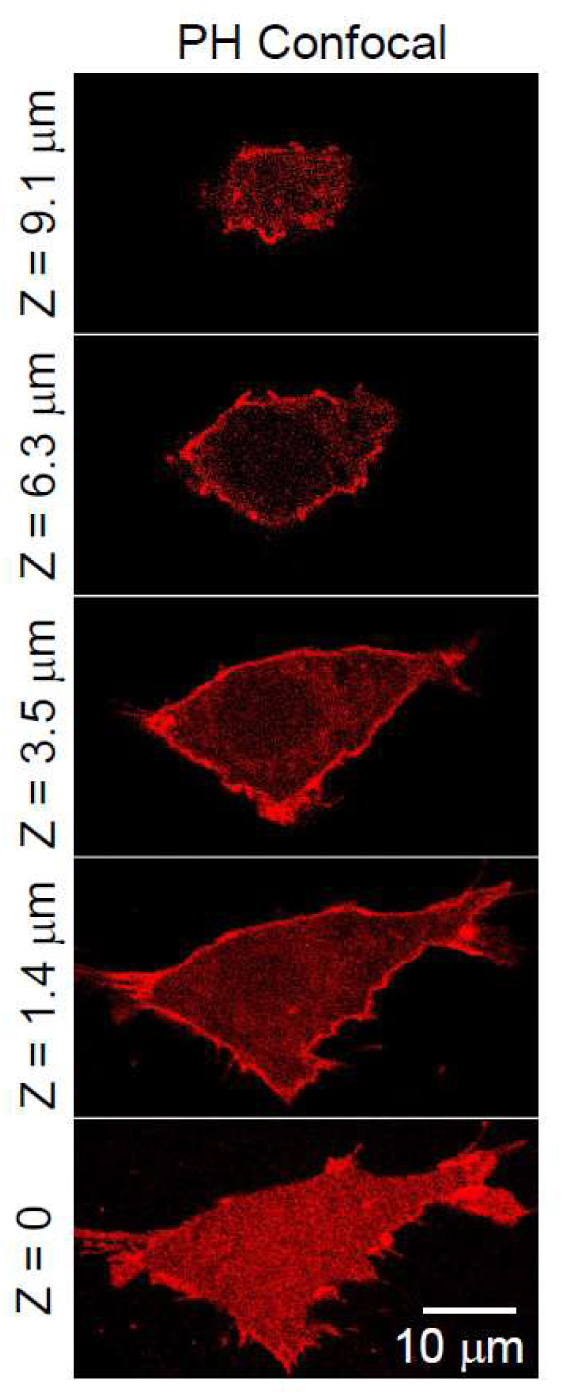
Sample images of the cell plasma membrane labelled with PH_SNAP_. Confocal XY-plane images of PH_SNAP_ at various Z-axis locations as labelled. A series of XY-plane images along the Z-axis every 700 nm were used to estimate the cell-surface area.

**Figure S9.**
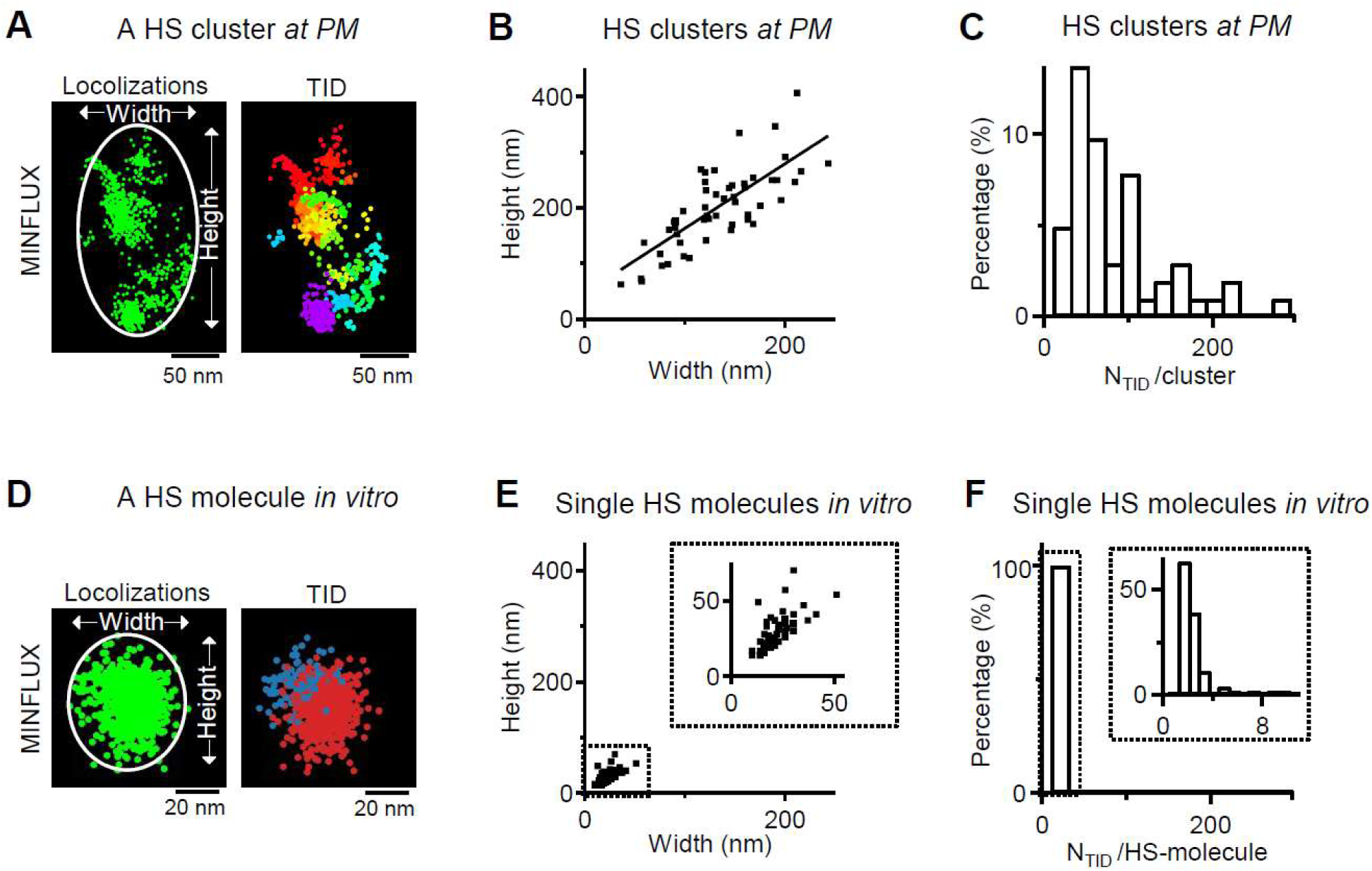
MINFLUX images of HS clusters at cell surface and single HS molecules in vitro. **A, Left:** MINFLUX image of a cluster of immunolabelled HS at the plasma membrane (PM). Height and width were measured by fitting the localizations in the cluster with an ellipse (Image J). **Right:** Same data as in the left, but plotted with localization trace IDs (TID) labeled in different colors. **B,** Height plotted versus width of HS clusters at the plasma membrane (53 clusters, 9 cells). The line is a linear regression fit (correlation coefficient: 0.8). **C,** Distribution of the number of TIDs per HS cluster (N_TID_/cluster, 53 clusters, 9 cells). **D, Left:** MINFLUX localization image of a single HS molecule in vitro. Purified HS was plated on the coverslip and labeled with antibodies. Height and width were measured by fitting the localizations with an ellipse (Image J). **Right:** Same data as in the left, but plotted with localization TIDs labeled in different colors. **E,** Height plotted versus width of single HS-molecule localizations in vitro (67 HS-molecules). The dotted box is enlarged in the inset. **F,** Distribution of the number of TIDs per single HS-molecule (N_TID_/HS-molecule). The dotted box is enlarged in the inset.

**Figure S10.**
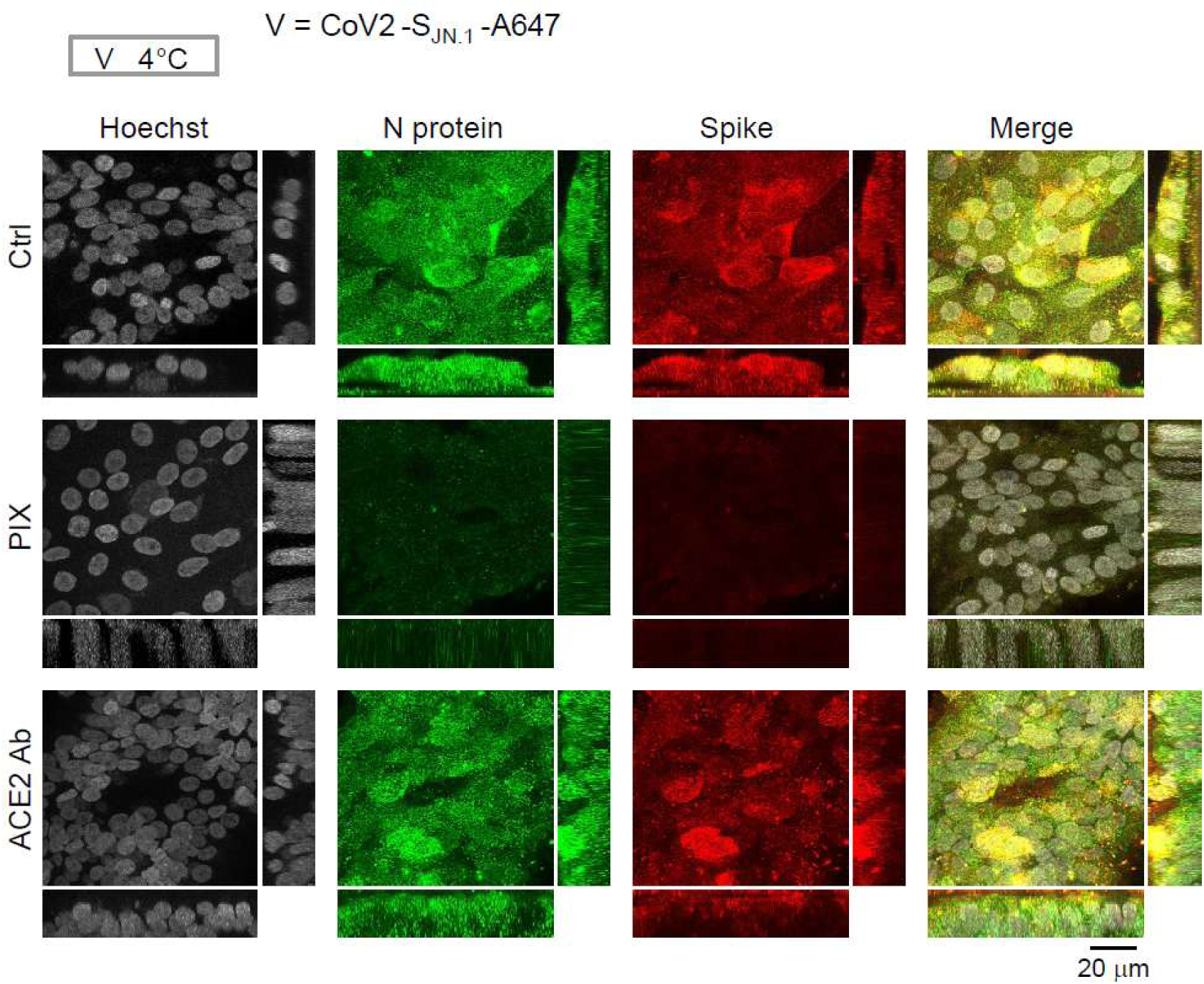
PIX, but not ACE2 antibody, inhibits CoV2-S_JN.1_ attachment in human primary small airway epithelial cells in air-liquid interface (ALI) format. Sampled confocal images of Hoechst (gray, labeling cell nucleus), antibody-labeled Nucleocapsid protein (N Protein of CoV2-S_JN.1_, green) and Spike protein (anti-S2P6 of CoV2-S_JN.1_, red) in human primary small airway epithelial cells in air-liquid interface (ALI) format; ALI cells in control (Ctrl), pre-treated with Pixantrone (PIX), or pre-treated with Anti-ACE2 antibody (ACE2 Ab) were incubated with CoV2-S_JN.1_ for 1 h at 4°C (for viral cell-surface attachment, protocol shown at the top). Images were projection images. XZ- and YZ-plane images are also shown on the bottom and right, respectively.

